# Structural Basis for Regulated Assembly of the Mitochondrial Fission GTPase Drp1

**DOI:** 10.1101/2023.06.22.546081

**Authors:** Kristy Rochon, Brianna L. Bauer, Nathaniel A. Roethler, Yuli Buckley, Chih-Chia Su, Wei Huang, Rajesh Ramachandran, Maria S. K. Stoll, Edward W. Yu, Derek J. Taylor, Jason A. Mears

## Abstract

Mitochondrial fission is crucial for distributing cellular energy throughout cells and for isolating damaged regions of the organelle that are targeted for degradation. This multistep process is initiated by the enhanced recruitment and oligomerization of dynamin-related protein 1 (Drp1) at the surface of mitochondria. As such, Drp1 is essential for inducing mitochondrial division in mammalian cells, and homologous proteins are found in all eukaryotes. *De novo* missense mutations in the Drp1 gene, DNM1L, are associated with severe neurodevelopmental diseases in patients, and no effective treatments are available. As a member of the dynamin superfamily of proteins (DSPs), controlled Drp1 self-assembly into large helical polymers stimulates its GTPase activity to promote membrane constriction. Still, little is known about the regulatory mechanisms that determine when and where Drp1 self-assembles, and proper mitochondrial dynamics requires correct spatial and temporal assembly of the fission machinery. Here we present a cryo-EM structure of a full-length, native Drp1 dimer in an auto-inhibited state. This dimer reveals two key conformational rearrangements that must be unlocked through intermolecular interactions to achieve the assembly competent state previously observed in crystal and filament structures. Specifically, the G domain is closed against the stalk domain and occludes intermolecular interactions necessary for self-assembly beyond a dimer. Similarly, adjacent stalks in the dimer form a more continuous interface that further occludes conserved intermolecular contact sites. This structural insight provides a novel mechanism for regulated self-assembly of the mitochondrial fission machinery.

## Background

Dynamin superfamily proteins (DSPs) are a group of large, multi-domain GTPase proteins with a conserved catalytic domain and a stalk capable of driving self-assembly of oligomers and helical polymers. Found in bacteria and eukaryotic cells, their primary function is membrane remodeling, as distinct family members play important roles in vesicle budding and organelle fission and fusion. Given their important and varied roles, structural studies have sought to characterize DSP domain organization and contribution to membrane remodeling. In part, this was pursued using cryo-EM to examine DSP interactions on membrane templates or in complex with partner proteins in helical assemblies or filaments^1–4^. Through considerable effort, crystal structures were determined for individual DSP domains^5–7^ and then for full-length proteins^8, 9^. DSPs form insoluble assemblies at the higher concentrations required for crystallization, so several mutations were introduced to improve solubility^9–11^.

Dynamin, the DSP founding member, has been studied for nearly four decades and there are still fundamental questions regarding the mode of assembly from cytosolic protein to the larger contractile machinery. All DSPs have two common domains, a GTPase or G domain, where GTP binding and hydrolysis occurs and the stalk, comprised of the middle domain and GTPase effector domain (GED), which together drive self-assembly (Fig. 1a). Fission DSPs include an additional bundle signaling element (BSE) connecting the stalk and G domain. The BSE changes conformation in response to nucleotide state to drive a powerstroke in dynamin that induces constriction of the helical polymer^12^. Most DSPs have additional domains to serve specialized roles within the cell. Dynamin-related protein 1 (Drp1), the master regulator of mitochondrial fission, has a unique intervening sequence adjacent to the stalk called the variable domain (VD), composed of 136 intrinsically disordered residues that confer lipid sensing^13^. This region also regulates Drp1 assembly properties^14^, and the disordered nature is necessary for function but likely prevents crystal formation and was deleted in the crystallographic studies.

**Fig. 1:**
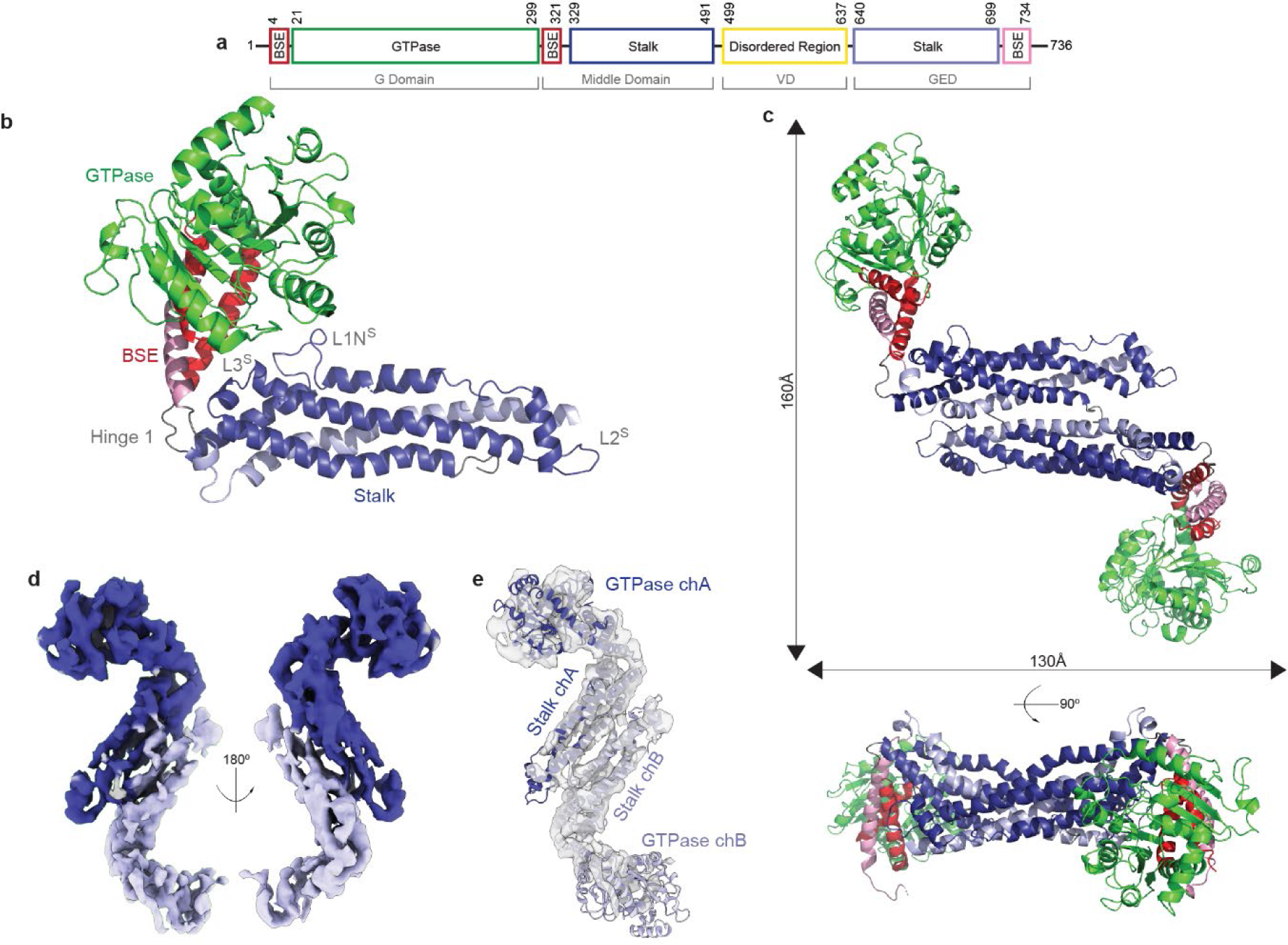
Native cryo-EM Structure of Drp1. **a**, Domains (Bundling Signaling Element, GTPase, Stalk, and Disordered Region) and their corresponding residues were identified previously in human Drp1. ^9^ The residue numbers represent the architecture of Drp1 isoform 1 (Drp1-1). The DSP domains are labeled in gray. **b**, Chain A (chA) of solution structure resolved using single particle, cryo-EM. The GTPase domain (green), BSE (red) and stalk region (blue) are indicated. The variable domain (VD) is not shown as this was not resolved. **c**, Top down and side view of the dimer of solution structure resolved using single particle cryo-EM. **d**, Density of the native cryo-EM dimer of Drp1 (dark blue = chain A, light blue = chain B). **e**, Docked structure within cryo-EM density after refinement.

### Identification of WT Drp1 Dimer Structure

Drp1 has been shown to exist as a mixture of dimers and tetramers in solution in a concentration dependent manner, and the dimer state represents the core unit of the larger helical machinery^15^. Dimers were isolated for study by diluting the protein to concentrations that would be enriched for dimers (600 nM, Extended Data Fig. 3d). Using cryo-EM, the structure of a native dimer of WT human Drp1 was resolved to a global resolution of 5.97 Å (Table 1, Extended Data Fig. 1). Four conformations were identified with significant conformational heterogeneity conferred primarily through GTPase motions that highlight variability in the position of this domain relative to the stalk. Overall, the architecture of the dimer presents a compact organization of the GTPase and stalk domains (Fig. 1b, Supplementary Videos 1 and 2), resulting in a ∼100Å decrease in length of the solution dimer when compared to the length of the crystal structure dimer with the VD deletion and GPRP401-404AAAA mutation (Extended Data Fig. 1e). Importantly, this conformation is incompatible with assembly into larger helical complexes as stalk interfaces that mediate assembly of DSP helical structures are occluded. The compaction of Drp1 dimers, as compared to the extended state described in the crystal structure, is achieved through hinge motions. Specifically, hinge 1, comprised of two disordered loops that connect the BSE to the distal end of the stalk, adopts a unique conformation that places the GTPase domain near its own stalk. Separately, a pivot between adjacent stalks in the homodimer increases the number of intermolecular contacts as compared to those in the crystal structure.

The largest rearrangement when comparing the solution Drp1 to previous structures was observed in hinge 1, connecting the BSE and stalk domains. A local resolution of 5.5 Å was observed in this region (Extended Data Fig. 1c), allowing flexible fitting of the crystal structure to the EM density. In comparison with the crystal structure, the fit revealed that the G domain is capable of significant rearrangement to bring the GTPase domain helix α2^G^ adjacent to loop L1N^S^ in the stalk (Fig. 1b). This loop is important for Drp1 assembly, as mutations in this conserved region limit Drp1 and other DSPs ability to form dimers^4, 15–17^. The compaction of the GTPase domain against this loop would prevent assembly beyond a dimer, representing a conformational change compared to helical and filament assemblies that exhibit an extended G domain conformation to expose conserved self-assembly interfaces.

The residues that form the discrete dimer interface identified by the Daumke group remain at the core of this dimer conformation^9^; however, the density reveals additional burial of surface area between adjacent stalks in this region. Additional residue contacts are formed through stalk motions that close the angle between the monomers. The orientation of head and stalk is generally heterogenous and asymmetric, suggesting that the hinge motion is dynamic in the solution state of the Drp1 homodimer (Extended Data Fig. 1). The major difference in the different dimer conformations can be attributed to a “pivoting” motion around the fulcrum of the interface that results in altered angles between adjacent stalks., However, compared to the crystal structure, the solution dimer conformation is more closed, further obscuring potential stalk interfaces. The previous structures in the crystal lattice or assembled helical polymers likely represent an “open” conformation where the core dimer interface is maintained but the peripheral stalk regions are exposed, removing auto-inhibitory interactions that limit assembly in these regions.

### BSE Lock through Loop L3^S^

The G domain moves 79 Å with a 67° rotation and a 61° twist (Fig. 2a and b). This results in a “locked” conformation of the G domain against the distal end of the stalk, mediated through interactions between the BSE and loop L3^S^ (Fig. 2c, Supplementary Video 3). Absent side chain resolution to identify critical contacts stabilizing this conformation, mutagenesis was pursued to further examine the role of this loop. L3^s^ was not fully resolved in the crystal structure; however, a Drp1-MiD49 filament structure^4^ identified a small helical segment in the middle of the loop at residues 452-456 QELLR, and AlphaFold predicted a helix in this region as well^4, 18^. A comparison of available DSP structures reveals that mitochondrial fission DSP loops have additional residues (448-451 NYST) when compared with other DSPs (Extended Data Fig. 2c-e). This additional sequence may contribute to the BSE lock, and a similar conformational rearrangement was observed by the Low group in a crystal structure tetramer of *Cyanidioschyzon merolae* Dmn1 (cmDmn1) ^19^, the red algae mitochondrial fission dynamin (Extended Data Fig. 3f). Dynamin has a shorter loop with a more extended α2^S^ helix, so it remains unclear whether the BSE lock is conserved in all DSPs. If this feature is unique to mitochondrial fission DSPs, this region would affect the wide range of BSE motions depending on its activation state (Extended Data Fig. 2g).

**Fig. 2:**
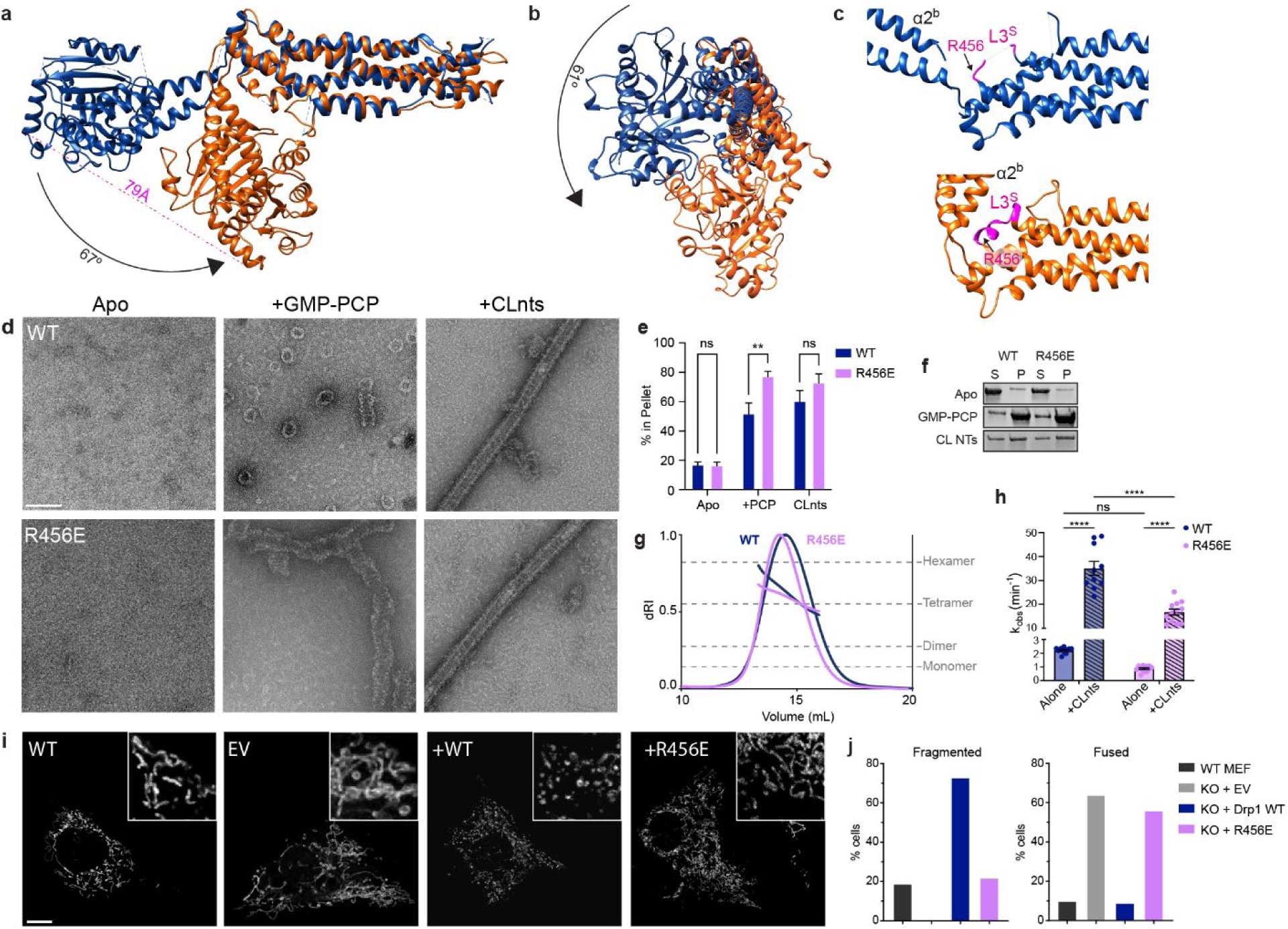
The BSE lock. **a-b**, Comparing the crystal structure (AlphaFold, blue) to the native cryo-EM structure (orange) based on overlay of the stalk region. **c**, Loop 3 (L3^s^) interacts with BSE helix α2^b^ and contains R456. **d**, Negative stain electron microscopy characterization of WT and R456E Drp1 in the presence and absence of non-hydrolysable GMP-PCP and CL-containing nanotubes (CLnts). *Scale bar = 100 nm* **e-f**, Sedimentation assays quantify changes in polymerization. Bars show the percent of protein detected in the pellet. Error bars are percent error. A two-tail t test was used to determine statistical significance. N-values WT=9, R456E=8. *\*\* P = 0.006*. **g**, SEC-MALS evaluation of Drp1 WT and Drp1 R456E. **h**, GTPase activity was determined for WT and R456 Drp1 alone and stimulated with CLnts. N-values WT (apo)=12, WT (nts)=10, R456E (apo and nts)=12. \*\*\*\**P<0.0001.* **i**, WT MEFs were compared to Drp1 knock-out MEFs which were transfected with empty vector (EV), Drp1 WT, and Drp1 R456E. MitoTracker Orange was used assess mitochondrial morphology. *Scale bar 5 µm, inset 10 µm.* **j**, Mitochondrial morphology was quantified using a blinded assessment described in the methods section. N-values WT MEF=40, EV=42, KO+WT=27, R456E=37.

To determine the effect of this small helical segment in L3^S^ on the activity of Drp1, a charge reversal was introduced through a charge reversal mutation, R456E, to disrupt this lock and promote the opening of the BSE away from the stalk. Previous work in the Mears lab looked at mutations in the N-terminal side of L3^s^ by mutating the glutamine and glutamic acid to a pair of lysines, similar to those found in the dynamin sequence. No difference was observed in the QE452-453KK mutation when compared to WT, as the GTPase and assembly activities were maintained with this change. The GMP-PCP-bound cryo-EM structure of Drp1^4^ reveals an extension of the G domain away from L3^s^ (Extended Data Fig. 2g), showing the range of BSE extension possible in this region of Drp1. Indeed, when GMP-PCP was added to Drp1 R456E, the protein had an increased propensity to assemble into spiral polymers compared to WT, consistent with this mutation opening the auto-inhibited lock(Fig. 2d). In agreement, 77% of the R456E protein sedimented in the presence of GMP-PCP as compared to only 52% for WT protein, further suggesting that R456E favors assembly in a GTP-bound state (Fig. 2e). The decoration on cardiolipin-containing nanotubes (CLnts) observed by negative stain was similar to WT, so the organization of the polymer does not appear to be affected by this mutation; rather, the mutation most likely results in a protein conformation that is more poised to assemble in the right condition, including lipid interactions with the VD and nucleotide binding. Importantly, there was no appreciable aggregation with the WT or R456E (Fig. 2e and f), and there was no shift in its solution multimeric state when assessed using size-exclusion chromatography coupled to multi-angle light scattering (SEC-MALS, Fig. 2g). Together, these data show the BSE lock did not impact the solution multimer formation. Instead, the charge reversal in L3^S^ weakens this self-regulatory BSE lock, priming the protein for assembly. The lipid sensing regions of the variable domain and the stalk interfaces required to build the helical assembly were not perturbed.

To complement the assembly assay, enzymatic activity was measured (Fig. 2h) and R456E (0.32 min^−1^) exhibited a five-fold decrease in basal activity compared WT (1.7 min^−1^). This change likely reflects conformational differences in the protein since the multimer state in solution is not altered. The stimulated activity of R456E in the presence of CLnts (16.8 min^−1^) was only two-fold lower compared to WT (35.1 min^−1^), so the magnitude of stimulation when compared to basal rates was greater for R456E (∼53-fold) than WT (∼21-fold). This highlights the greater assembly-stimulated activity of R456E when compared to WT, similar to the results in the presence of GMP-PCP. Again, this mutant is more potent at assembly, and the lipid-bound structures look indistinguishable (Fig. 2d), though we cannot discount small differences that affect this stimulation.

To test the effect of the mutation on mitochondrial fission in cells, we transfected Drp1 knock-out (KO) mouse embryotic fibroblasts (MEFs) with WT Drp1 and R456E (Fig. 2i). When transfected with WT Drp1, cellular mitochondrial networks were mostly fragmented due to the over-expression of the fission protein. Cells transfected with R456E were observed to have fused mitochondria, similar to the KO treated with an empty vector (EV) control. Additionally, the R456E mutant protein in these transfected cells organized in aggregated puncta and did not exhibit a diffuse signal observed with the WT Drp1 vector (Extended Data Fig. 2h). This likely represents premature assembly of the R456E mutant in cells, suggesting that this BSE lock is critical to sustain an auto-inhibited state that prevents premature assembly of Drp1 polymers. Only when this autoinhibited state is relieved, or unlocked, does the WT protein become primed to form a helical assembly around the outer mitochondrial membrane.

### A Flexible Dimer Interface

The dimer interface that orients the monomers relative to one another was found to have a large amount of heterogeneity within the conformations and among available structural data. The crystal structure exhibits a discrete dimer interface with a more acute lateral angle measuring 85° (Fig. 3a). The solution structure presented here suggests that removing the VD and introducing the poly-A mutation within L2^s^ near the membrane proximal interface (previously labeled interface 3) yielded an open conformation more amenable to crystallization. These changes also disrupted key regulatory regions, leading to an active conformation, primed for assembly, and the stalk orientations are consistent with previously reported DSP helical conformations. Within the cell, interactions with lipids, ER contact sites, post-translational modification(s), and partner proteins interactions could all promote an open state alone or in concert with one another. In the solution (i.e. “closed”) state, L2^s^ is juxtaposed to α1N^s^, and this interaction requires a conformation with an obtuse angle between adjacent stalks (ranging from 103-145°) to form a more continuous interface (Extended Data Fig. 3a). In order to identify intra-monomer conformational rearrangements, the four helices comprising the stalk of the solution structure were aligned with the crystal structure helices (Fig. 3b). Loops within α1 confer a large degree of flexibility, and this is evident when comparing the different chains of the crystal structure (Extended Data Fig. 3c). Helices 2 and 4 were relatively unchanged, and Helix 3 had a shift in alignment at the C-terminal end due to flexion in the middle of the helix. A conserved tyrosine (Y493) was found to bookend the dimer interface based on optimal placement of this helix in the density (Extended Data Fig. 3b, Supplementary Video 4). To alter the interactions at this position, mutagenesis substituted a smaller alanine in place of the bulkier tyrosine. As evidenced by negative-stain EM analysis, this mutation showed a decreased ability to form spirals in the presence of GMP-PCP and exhibits a decreased ability to decorate CLnts when compared to WT (Fig. 3d). This finding was further quantified using a sedimentation assay. In apo conditions, both WT and Y493A resulted in similar levels of protein found in the pellet (17% and 20% respectively). No assembly of Y493A was detected when GMP-PCP was added to induce spiral formation. We found 19% sediments for Y493A with GMP-PCP representing no change compared to average sedimentation into the pellet under apo conditions, while WT Drp1 had a three-fold increase in pelleted protein (52%). A decrease was observed in the ability of Y493A to decorate CLnts (42% of Y493A was found in the pellet compared to 60% for WT), confirming that this mutant prevents assembly even though lipid binding is preserved since the VD is unchanged. There was no significant change in sedimentation when comparing the WT and Y493A Drp1 in the absence of assembly inducers (Fig. 3e and f), so SEC-MALS was used to assess the multimer state in solution. Y493A was found to exist in equilibrium between dimers and monomers and was not able to form larger multimers observed with WT Drp1 at the same concentration (Fig. 3g). To confirm this difference and knowing oligomer state for Drp1 is concentration dependent, mass photometry was used at a lower concentration (100 nM), and Y493A was found to exist largely as a monomer while WT protein was mostly dimeric (Extended Data Fig. 3d).

**Fig. 3:**
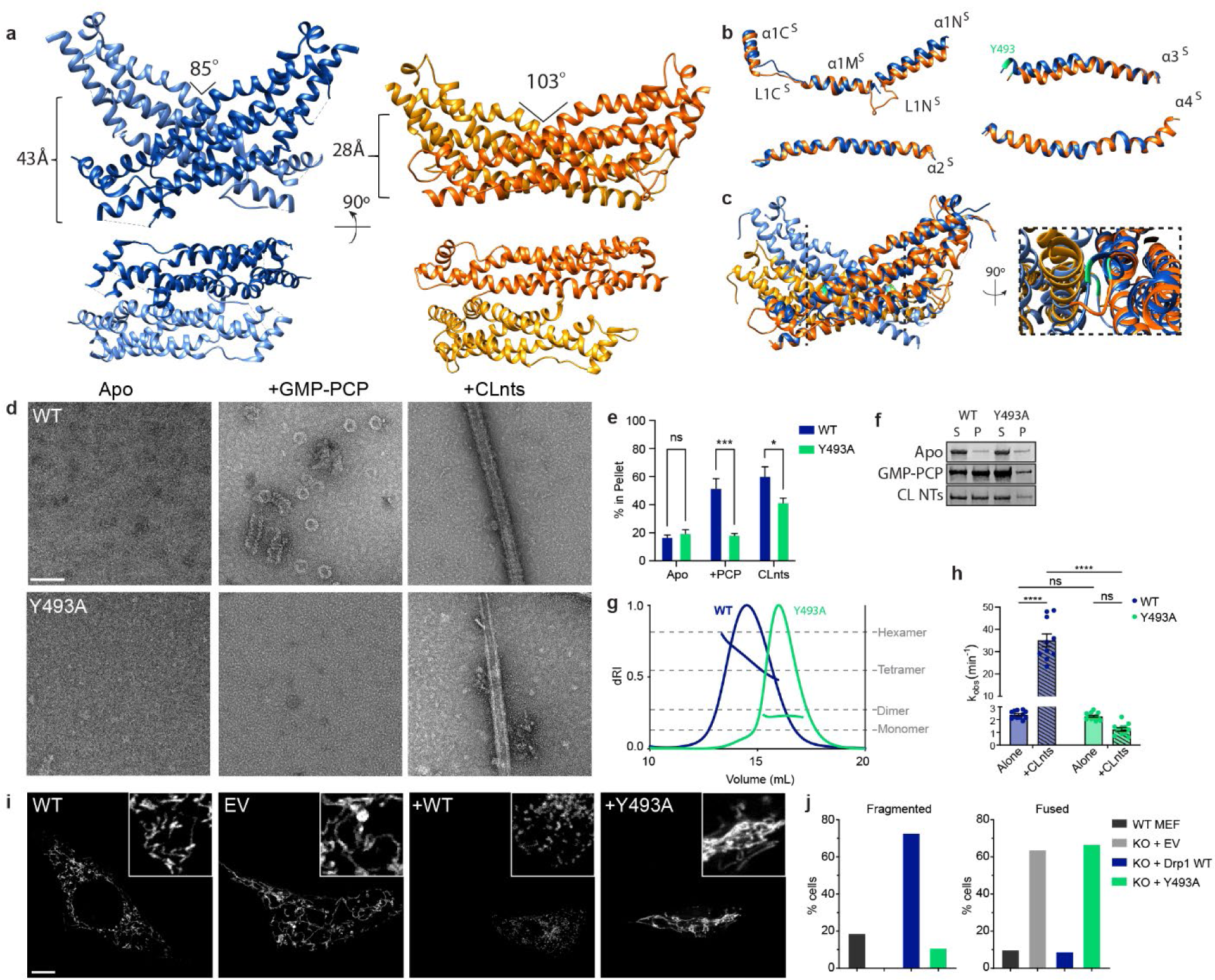
Native cryo-EM Structure Dimer Interface. **a**, Comparison of the stalk orientations in the crystal structure dimer interface, (PDB code: 4BEJ/AlphaFold, dark blue chain A, light blue chain B) and the native cryo-EM structure (dark orange chain A, light orange chain B). **b**, Alignment of each helix comprising the stalk domain (PDB code: 4BEJ blue, solution structure orange). Y493 is highlighted in light green. **c**, Alignment of crystal structure dimer and native cryo-EM structure stalks, reference chain is chain A. Zoomed in view, Y493 in light green. **d**, Negative stain electron microscopy characterization of WT and Y493A Drp1 in the presence and absence of non-hydrolysable GMP-PCP and CL-containing nanotubes (CLnts). *Scale bar = 100nm* **e-f**, Sedimentation assays were used to quantify changes in polymerization. Bars show the percent of protein in the pellet. Error bars are percent error. A two-tail t test was used to determine statistical significance. N-values WT=9, R456E=8.*\*\*\*P=0.0005; *P=0.04*. **g**, SEC-MALS evaluation of Drp1 WT and Drp1 Y493A is presented. **h**, GTPase activity was assessed for WT and Y493A Drp1 alone and stimulated with CLnts. N-values WT (apo)=12, WT (nts)=10, Y493A (apo)=12, Y493A (nts)=11. \*\*\*\**P<0.0001.* **i**, WT MEFs were compared to Drp1 KO MEFs transfected with empty vector (EV), Drp1 WT, and Drp1 Y493A. MitoTracker Orange was used to assess mitochondrial morphology. *Scale bar 5 µm. Inset 10 µm*. **j**, Mitochondrial morphology was quantified using a blinded assessment described in the methods section. N-values WT MEF=40, EV=42, KO+WT=27, Y493A=23.

The GTPase activity was assessed for Y493A and WT Drp1 (Fig. 3h), and the basal rates were comparable (2.3 min^−1^ for Y493A versus 2.4 min^−1^); however, CLnts stimulation was only observed with the WT protein (35 min^−1^), while Y493A was not stimulated (1.2 min^−1^). Therefore, Y493A maintained basal GTPase activity, but it was incapable of functional assembly.

Transfecting Drp1 KO MEF cells with Y493A saw no change in the interconnected, fused mitochondrial morphology when compared to the empty vector control. Conversely, WT Drp1 transfection resulted in a fragmented mitochondrial network (Fig. 3i and 3j). This observation is consistent with the destabilization of the continuous dimer interface by the Y493A mutation that was introduced, which prevents functional assembly and limits mitochondrial fission in cells. Therefore, the Y493 residue is critical for stabilizing interface 2, acting as a “shock absorber” that accommodates flexibility from a continuous interface to a more discreet intermolecular interaction required for spiral and helical oligomerization.

## Discussion

This cryo-EM solution structure of WT Drp1 demonstrates a previously unappreciated inactive dimer conformation through additional self-regulatory interactions. These new insights will be important for developing new selective inhibitors of Drp1. To date, efforts to regulate Drp1 have been developed based on the field’s understanding of Drp1 already in an open and active conformation and have failed to produce a reliable inhibitor targeting Drp1 directly. With other DSPs, conformational changes were observed in the BSE hinge in response to nucleotide binding and partner protein interactions^4, 12^. These motions were proposed to alter stalk interactions to mediate constriction of the helical lattice. The solution structures of Drp1 demonstrate additional crosstalk between the G domain and stalk regions that offer a new role for accessory protein and lipid interactions in relieving autoinhibitory conformations (Figure 4). In the absence of an activating stimulus, the BSE lock is engaged to prevent premature assembly. This conformation is associated with increased stalk interactions, forming a more continuous dimer interface. Together, these changes compared to the crystal structure represent key regulatory motions that open the dimer preceding functional assembly.

**Figure 4:**
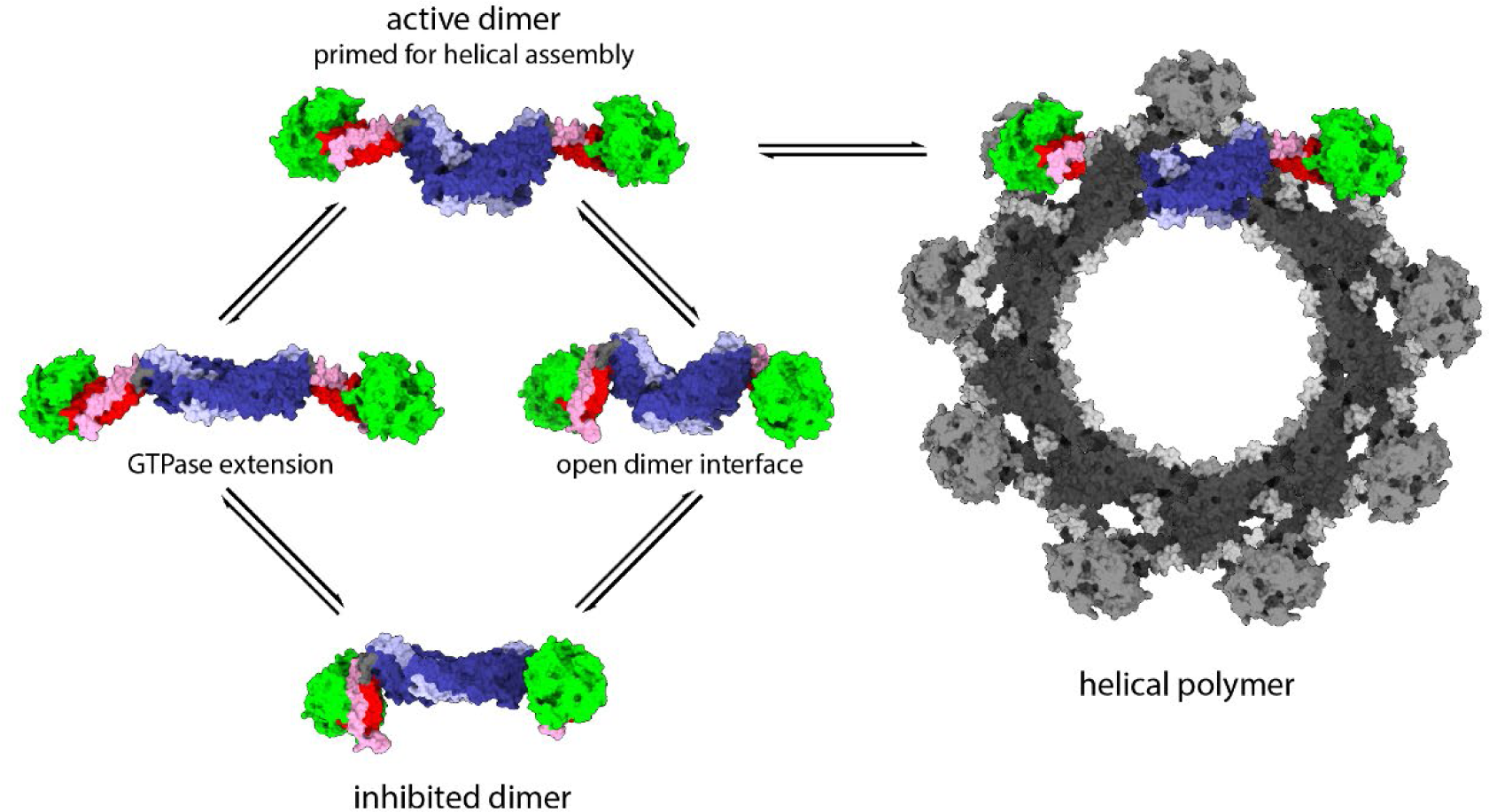
New Model for Drp1 Regulation. The native cryo-EM structure (bottom left) is an autoinhibited dimer. The BSE lock and the closed dimer interface are maintained by weak intramolecular reactions within the dimer. Intermolecular interactions between Drp1 and partner proteins, organelle contact sites, and/or lipid interactions relieve inhibition to promote an assembly primed conformation. Helical assembly requires both an open dimer interface and an unlocked, extended G Domain. GTPase domain in green, stalk in dark blue, GED in light blue, BSE in red (N-terminal) and Pink (C-terminal). Variable domain not shown.

A nucleotide-bound, cryo-EM structure of a Drp1-MiD49 co-polymer was solved with full-length Drp1 in an open conformation^4^, and it also exhibited a more extended BSE due to partner protein binding interactions at stalk interfaces (Extended Data Fig. 3g). When comparing the solution structure with the MiD49-bound filament, the “closed” BSE lock is incompatible with two MiD49 interfaces (Extended Data Fig. 3h and i). One proposed MiD49 interface is positioned at loop 1N^s^ (Extended Data Fig. 3j), which would require that the G Domain extend to avoid steric clashes, consistent with altered hydrolysis, and be available for G-G interactions that stabilized inter-rung interactions in the helical lattice. Since MiD49 is anchored at the surface of mitochondria, this interaction would favor opening of Drp1 to promote assembly at these defined membrane sites.

Nucleotide interactions could also facilitate opening of hinge 1 through motions in the adjacent BSE hinge, and this model is consistent with nucleotide-induced formation of spiral structures observed when Drp1 is incubated with non-hydrolysable GTP analogs. It remains unclear whether the BSE hinge responds to nucleotide binding, since nucleotide binding does not lead to constriction like it does for dynamin. Rather, nucleotide binding likely augments release of the BSE lock to promote assembly. Thus, protein or nucleotide interactions change the G Domain conformation to inform conformational rearrangements in the stalk that permit assembly of the helical fission complex.

A separate MiD49 interface at loop 2^s^ is likely regulated by interactions with the VD. In the Drp1-MiD 49 structure, the binding of the partner protein would occlude VD interactions as previously suggested^13^. In the solution structure, VD and L2^s^ interactions stabilize stalk interactions and yield a more continuous dimer interface. Deletion of the VD in the crystal structure removed a large regulatory domain, and previous studies have shown that removing the VD predisposes Drp1 to self-assembly through interactions with partner proteins^20^ or when the protein concentration is elevated^15^. On the surface of mitochondria, VD engagement with these partners would promote opening of the dimer interface and expose key intermolecular interaction sites to drive helical formation.

In this new model for assembly of the mitochondrial fission machinery, locked G domains extend to promote helical assembly. In addition, the more continuous dimer interface observed in solution exhibits a wide range of flexibility within the dimer. This novel conformation occludes residues necessary for helical assembly, decreasing the allowable geometries needed for assembly of the fission machinery. We propose that these self-regulatory interactions exist in an equilibrium in solution, sampling various conformations. Assembly-limited mutants, often located in the L1^N^ loop, likely limit the range of conformational sampling and thereby prevent opening of the dimer to an active state. Conversely, stimulating interactions with specific lipid and partner proteins at the surface of mitochondria open the stalk configuration to promote intermolecular contacts between Drp1 dimers, which have been shown to be the core building block for the contractile helical polymer. Post-translational modifications could bias conformational sampling in the solution dimer to augment or prevent functional assembly. In this way, partner protein and lipid binding affinities reflect the accessibility of interaction sites on the Drp1 stalk that dictate further opening of the Drp1 dimer to build the mitochondrial fission apparatus in a manner that is spatiotemporally regulated.

## Materials and Methods

### Protein Constructs and Mutagenesis

Drp1 Isoform 1 (UniProt ID O00429-1) was cloned into the pCal-N-EK vector as described previously^20, 21^. Site-directed mutagenesis for Drp1 R456E and Y493A was completed to introduce each mutation into this construct individually using the QuikChange Lightning kit (Agilent) with primers from Integrated DNA Technologies (Coralville, IA).

### Protein Expression and Purification

All Drp1 constructs were expressed in BL21-(DE3) Star *Escherichia coli*. Cells were grown in LB containing 100 µg/mL ampicillin at 18°C with shaking at 200 rpm for 24 hours after induction with 1 mM isopropyl-1-thio-β-D-galactopyranoside (IPTG). Then, cells were harvested via centrifugation at 4,300 × g for 20 minutes at 4°C. The resulting pellet was resuspended in CalA Buffer (0.5 M L-Arginine pH 7.4, 0.3 M NaCl, 5 mM MgCl_2_, 2 mM CaCl_2_, 1 mM imidazole, 10 mM β-mercaptoethanol) with 1 mM Pefabloc-SC and 100 μg/mL lysozyme. Cells were lysed by sonication on ice. Next, the cell debris was pelleted via centrifugation at 150,000 × g for 1 hour at 4°C. First, the CBP-tagged Drp1 was purified by affinity chromatography using calmodulin agarose resin (Agilent) that had been pre-equilibrated with CalA Buffer. After the supernatant was loaded onto the column, the resin was washed with 25 column volumes of CalA Buffer. Next, 8 fractions of eluent were collected using 0.5 column volumes of CalB Buffer (0.5 M L-Arginine pH 7.4, 0.3 M NaCl, 2.5 mM EGTA, 10 mM β-mercaptoethanol). Protein-containing fractions were pooled and incubated with GST-tagged PreScission Protease (HRV-3C) overnight at 4°C to remove the CBP-tag. This solution was concentrated using a 30,000 molecular weight cut-off centrifugal filter (Amicon). This concentrated pool of Drp1 was further purified by size exclusion chromatography (SEC) with an ÄKTA Purifier FPLC (GE Healthcare) and a HiLoad 16/600 Superdex 200 Prep Grade column that had been pre-equilibrated with SEC Buffer (25 mM HEPES (KOH) pH 7.5, 0.15 M KCl, 5 mM MgCl_2_, 10 mM β-mercaptoethanol). All elution fractions containing Drp1 were pooled and concentrated once again, and glycerol (5% final) was added. The purified Drp1 was aliquoted, flash frozen in liquid nitrogen, and stored at −80°C until use.

### Cryo-EM Sample Preparation, Imaging, and Processing

Holey carbon grids (1.2/1.3, quantifoil) were coated with 0.2 mg/ml graphene oxide (GO) support film prepared in house. Drp1 was diluted to 0.05 mg/ml, incubated on the GO grid for 30 s, and then frozen using a FEI Vitrobot Mark IV with a blot force of 0 and a blot time of 2.0 s. Grids were imaged on a Titan Krios (300 keV) equipped with a K2 Summit camera at a magnification of 130,000x. Data were collected over 40 frames for 8 s at a nominal dose of 49 e/Å^2^ (6.11 e/p/s).

All data was processed using cryoSPARC, version 3^22^. Movies were motion corrected using MotionCor2^23^, and Patch CTF was used for contrast transfer function (CTF) estimation. Initial particles were picked manually (2,054) and then used to train TOPAZ^24^. Initial processing selected a subset of junk particles and good particles. Those were used to create to reference densities. All particles were then sorted using these references. 3D refinement was completed using the Nonuniform refinement job in cryoSPARC^25^. Local resolution was determined using the Local Resolution job in cryoSPARC and visualized with Chimera^26^.

### Model Building, Flexible Fit, and Refinement

The starting model was built using the AlphaFold predicted monomer^18, 27^. Two monomer chains were aligned to the crystal structure’s dimer interface (4BEJ) ^9^. This dimer model was docked using rigid body docking in Chimera. Then in VMD, an all-atom model was generated using the Automatic PSF Builder in VMD^28^. Files were prepared for NAMD processing following previously described methods^29^. The four density maps were converted to an MDFF potential. Secondary restraints were applied using NAMD’s extrabonds feature and plugins to preserve secondary structure and to limit artifacts to chiral centers and cis peptide bonds. The simulations used a GScale was set to 0.3, temperature was set to 300, and the number of timesteps (numsteps) was 1,000,000 – 1,500,000. A minimization step was run with a GScale was set to 10, temperature was set to 300, and minimize steps (minsteps) was set to 2000. The model was then minimized and refined using Phenix Real-space refinement^30, 31^ to fix clashes and outliers.

### Negative-Stain Electron Microscopy

All samples were added to 400 mesh copper grids with a formvar carbon film (EMS FCF400-CU) and stained using 2% uranyl acetate. Each grid was made using 2 μM protein and either 1 mM GMP-PCP or 150 μM lipid nanotubes as indicated. Sample images were acquired on a Tecnai TF20 electron microscope (FEI Co.) at 200 keV, respectively. The TF20 was equipped with TVIPS F-416 CMOS (4k × 4k) camera and images were acquired at a magnification of 30,000×.

### Malachite Green Colorimetric Assay

The basal GTPase activity of Drp1 was measured using a colorimetric assay to detect released phosphate, as described previously^20, 21^. Briefly, Drp1 (500 nM final) was diluted to 2.4X with Assembly Buffer (25 mM HEPES (KOH) pH 7.5, 150 mM KCl, 10 mM β-mercaptoethanol). To start the reaction, 3X GTP/MgCl_2_ (1 mM and 2 mM final, respectively) was added to the Drp1 with either 4X lipid (150 μM final) to calculate the lipid-stimulated rates or only Assembly Buffer to calculate the rate for the protein alone in solution. The reaction was carried out at 37°C. At the chosen time points, a sample aliquot was taken and quickly added to EDTA (100 mM final) to stop the reaction. The time points used to calculate the rate for the protein alone in solution were 5, 10, 20, 40, and 60 min. For the lipid-stimulated rates the time points were 2, 4, 6, 8, and 10 min for all samples except Y493A + CLnts, which was instead measured using the longer 5, 10, 20, 40, 60 min time course. After collecting all time points, Malachite Green Reagent (1 mM malachite green carbinol, 10 mM ammonium molybdate tetrahydrate, 1 N HCl) was added to each sample, and the absorbance at 650 nm was measured using a VersaMax microplate reader (Molecular Devices).

### Lipid Nanotube Synthesis

All lipid nanotubes utilized here were comprised by 40% D-galactosyl-beta-1’-N-nervonyl-erythro-sphingosine (GC), 35% phosphatidylethanolamine (PE). 25% bovine heart cardiolipin (CL) molar fractions. All lipids were purchased from Avanti Polar Lipids, Inc. (Alabaster, AL). Lipids were added to a glass test tube and slowly dried to a thin film using nitrogen gas. The lipid film was then stored in a desiccator for at least one hour to ensure any trace solvent remaining was removed. Then the lipid film was rehydrated with a buffer (200 μL) containing 50 mM HEPES (KOH) pH 7.5 and 0.15 M KCl and heated in a 37°C water bath for ∼40 minutes with gentle vortexing every 10 minutes. With these volumes, the final lipid nanotube concentration was 2 mM. The lipid film was placed in a water bath sonicator for 30 seconds and the resulting nanotubes were stored on ice until use.

### Size Exclusion Chromatography with Multi-Angle Light Scattering (SEC-MALS)

SEC-MALS experiments were performed as before^20^. Briefly, 5 µM Drp1 was injected onto a Superose 6 10/300GL column in an ÄKTApure FPLC system (GE Healthcare) connected in line with DAWN Heleos-II 18-angle MALS and Optilab T-rEX differential refractive index (dRI) detectors from Wyatt Technology. Data were analyzed with ASTRA 7 software from Wyatt Technology.

### Sedimentation Assay

To quantify Drp1 oligomerization, a sedimentation assay was conducted similar to what has been described previously^32, 33^. Large oligomers formed by Drp1 samples, in the presence of ligands, were found in the pellet after a medium speed centrifugation. Specifically, protein was diluted in HEPES KCl buffer to 2 μM, and specified WT and mutant samples were incubated at room temperature with lipid nanotubes (150 μM) and/or GMPPCP (2 mM) for at least 60 minutes. The mixtures were then spun at 15,000 rpm for 10 min in a tabletop centrifuge (Eppendorf). The supernatant and pellet fractions were separated, collected, and immediately mixed with SDS-PAGE loading dye (Bio-Rad) and heated briefly at 100 °C. These samples were run on an SDS-PAGE gel and stained with an InstantBlue Coomassie dye (Expedeon). Gels were scanned using an Odyssey XF Imaging System (Li-Cor) and densitometry analysis was done using the Image Studio Lite Ver 5.2.

### Cell Lines and Transfection Protocol

WT and Drp1 KO Murine embryonic fibroblasts (MEFs) as previously described^34^ were cultured in complete media (DMEM with 10% FBS supplemented with Penicillin/Streptomycin, L-Glutamine, Insulin-Transferrin-Sodium Selenite Supplement (ITS), αFibroblast Growth Factor (FGF), and Uridine). Constructs were cloned into a pCMV-Myc vector. 25,000-50,000 cells were plated and transfection of specified constructs or empty vector (6-13 µg/nL) was performed with Lipofectamine 2000 (Thermo Fisher Scientific, 1166819) and Opti-MEM (Gipco, 31985) for 24 hours.

### Immunofluorescent Staining and Assessing Mitochondrial Morphology

Cells were plated on glass-bottom dishes and were stained with 250 nM of MitoTracker™ Orange CMTMRos (Invitrogen, M7510) for 30 min then fixed in 4% paraformaldehyde (Alfa Aesar, 43368) for 15 min and stained with DAPI in PBS 1:10,000 (Invitrogen, D1306) for 5 min. 3 washes were performed between each step to reduce background. Cells were permeabilized with 1% triton, blocked with 10% BSA, and incubated overnight with 1:250 c-Myc antibody (Santa Cruz, sc40) in 10% BSA. They were then washed with PBS 3 times, 5 minutes each wash and treated with 1:500 Alexa Fluor Plus 488, highly cross absorbed (Invitrogen, A32723TR) in 10% BSA. Then washed 3 times with PBS. Images acquired using a Leica SP8 Gated STED confocal Microscope in the Light Microscopy Imaging Facility at Case Western Reserve University. Blinded individuals independently categorized each cell as either fused, tubular, intermediate, or fragmented for randomized confocal images.

### Fluorescence Quantification

To determine cellular fluorescence of transfected cells, the corrected total cell fluorescence (CTCF) was determined: Integrated Density – (Area of Selected Cell X Mean Fluorescence of Background) using protocols described previously^35^. The anti-Myc channel was opened in ImageJ. Each cell was isolated using the freeform tool, excluding the void left by the nucleus and measured. An additional area (18-20 μm^2^) without signal was selected using the circle tool to measure the background.

### Statistics

Statistics were done in Prism GraphPad using a two-tail t-test. All measurements were taken from distinct samples as biological replicates.

## Data Availability

The 3D EM density and resulting structure have been deposited in EMDB (EMDB-40967) and PDB (8T1H).

## Author Contributions

K.R. and J.A.M. conceived and planned the experiments. K.R., B.L.B., N.A.R., Y.B., R.R., and M.S.K.S. carried out the experiments. C.S. and W.H. assisted with data processing and model validation. K.R. designed the figures and drafted the manuscript with supervision of J.A.M and input from all authors. R.R., E.W.Y., and D.J.T. contributed to the interpretation of results.

## Acknowledgements

We would like to acknowledge use of the (Leica SP8 confocal microscope) in the Light Microscopy Imaging Facility at Case Western Reserve University made available through the Office of Research Infrastructure (NIH-ORIP) Shared Instrumentation Grant S10OD016164.

We would like to acknowledge the West/Midwest Consortium for High-Resolution Cryo Electron Microscopy at the University of California, Los Angeles for use of their facilities for data collection made available through the cooperative agreements grant (NIH U24) GM116792.

K.R. discloses support for the research of this work from the National Institute of General Medical Sciences [F31 GM139324]. J.A.M. discloses support for the research and publication of this work from National Institute of General Medical Sciences [R01 GM125844].

Molecular graphics images were produced using the UCSF Chimera package from the Resource for Biocomputing, Visualization, and Informatics at the University of California, San Francisco (supported by NIH P41 RR-01081).

We would like to thank Harry Scott for his expertise in sample preparation and screening.

---

**Extended Data Fig 1:**
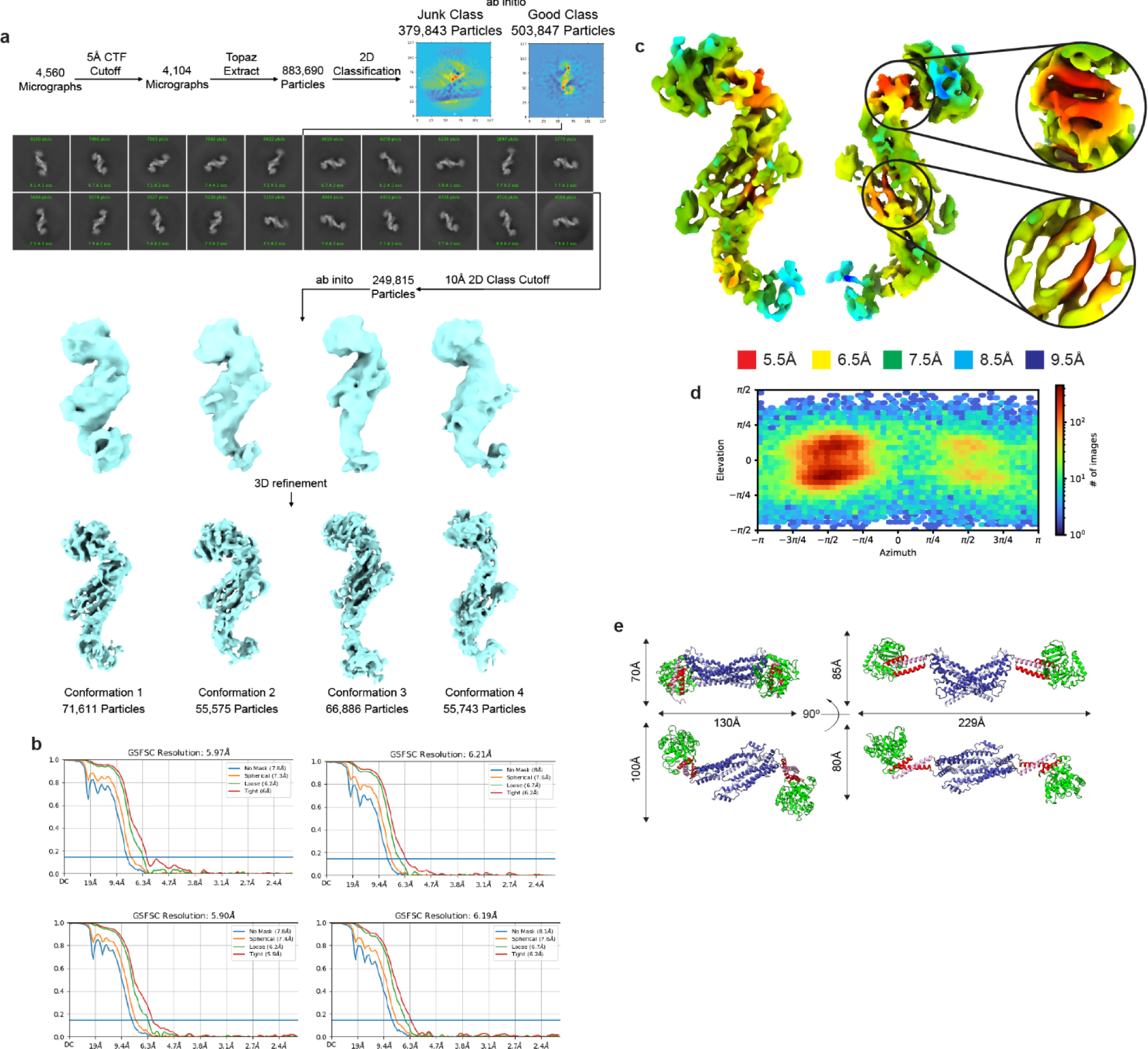
Single Particle Workflow. **a**, cryoSPARC preprocessing, classification, and refinement workflow using Topaz particle training. Four conformations were refined. **b**, Conformation 1 reported GSFSC is 5.97Å. Conformation 2 reported GSFSC is 6.21Å. Conformation 3 reported GSFSC is 5.90Å. Conformation 4 reported GSFSC is 6.19Å. **c**, xestimates in cryoSPARC of conformation 1 is highest within the dimer interface and at the BSE. **d**, The direction distribution of the particle stack confirms a preferred orientation, limiting the resolution from certain angles. **e**, The native cryo-EM structure (left) compared to crystal structure (4BEJ, right) modeled using the AlphaFold monomer and aligning two chains to the PDB 4BEJ dimer chains A and B.

**Extended Data Fig 2:**
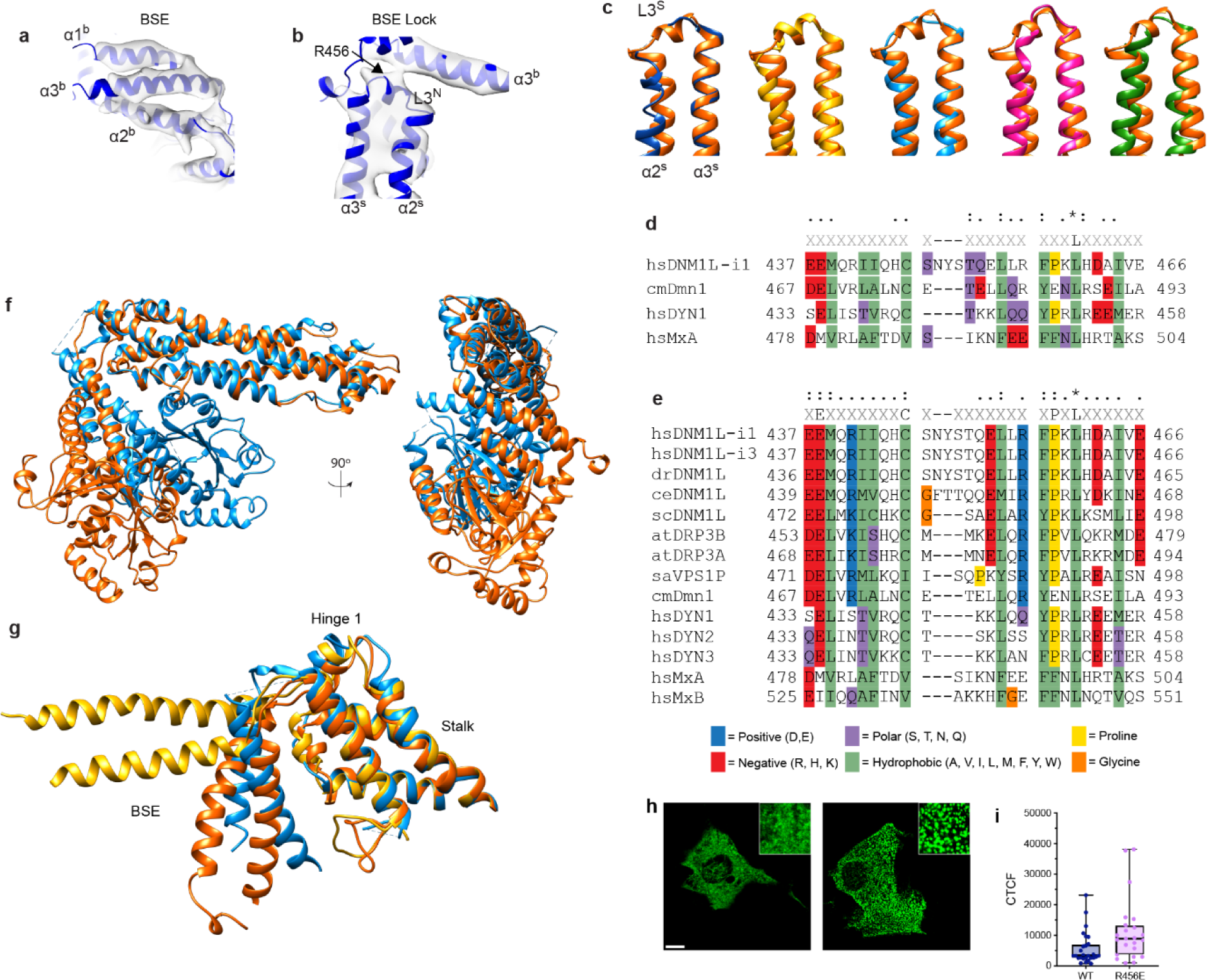
DSP structural comparison of BSE lock. **a**, All helices in the three-helix BSE fits within the density. **b**, The features contributing to the BSE lock against the stalk are observed within the density. R456 is found in the center of the lock. **c**, Selected DSP loop 3 comparison by aligning α3^s^ (Drp1 solution structure, orange; Drp1 crystal structure 4BEJ, dark blue; Drp1 cryo-EM filament with MID49 and GMP-PCP 5WP9, yellow; CmDnm1 crystal structure 6FGZ, light blue; Dyn1 cryoEM GTP-bound polymer 6DLV, pink; MxA crystal structure 3SZR, green). **d**, Sequence alignment of loop 3 of the DSPs above. - = no sequence or >15% consensus | X = Any amino acid, at least 15%, no consensus |. = > 35% < 75% |: = > 75% < 100% | * = 100%. **e**, Sequence alignment of loop 3 of all DSP aligned in Supplementary Data 1. **f**, Structural comparison of the G Domain position of Drp1 native cryo-EM structure (orange) and CmDnm1 (6FGZ, light blue). **g**, Structural comparison of BSE position of Drp1 solution structure (orange), CmDnm1 (6FGZ, light blue), and Drp1 nucleotide-bound filament (5WP9, yellow). **h**, Myc-tagged Drp1 in transfected Drp1 KO MEF cells. *Scale bar 5 µm, inset 5 µm.* **i**, Corrected Total Cell Fluorescence (CTCF) quantification for transfected cells shown in fig. 2.

**Extended Data Fig 3:**
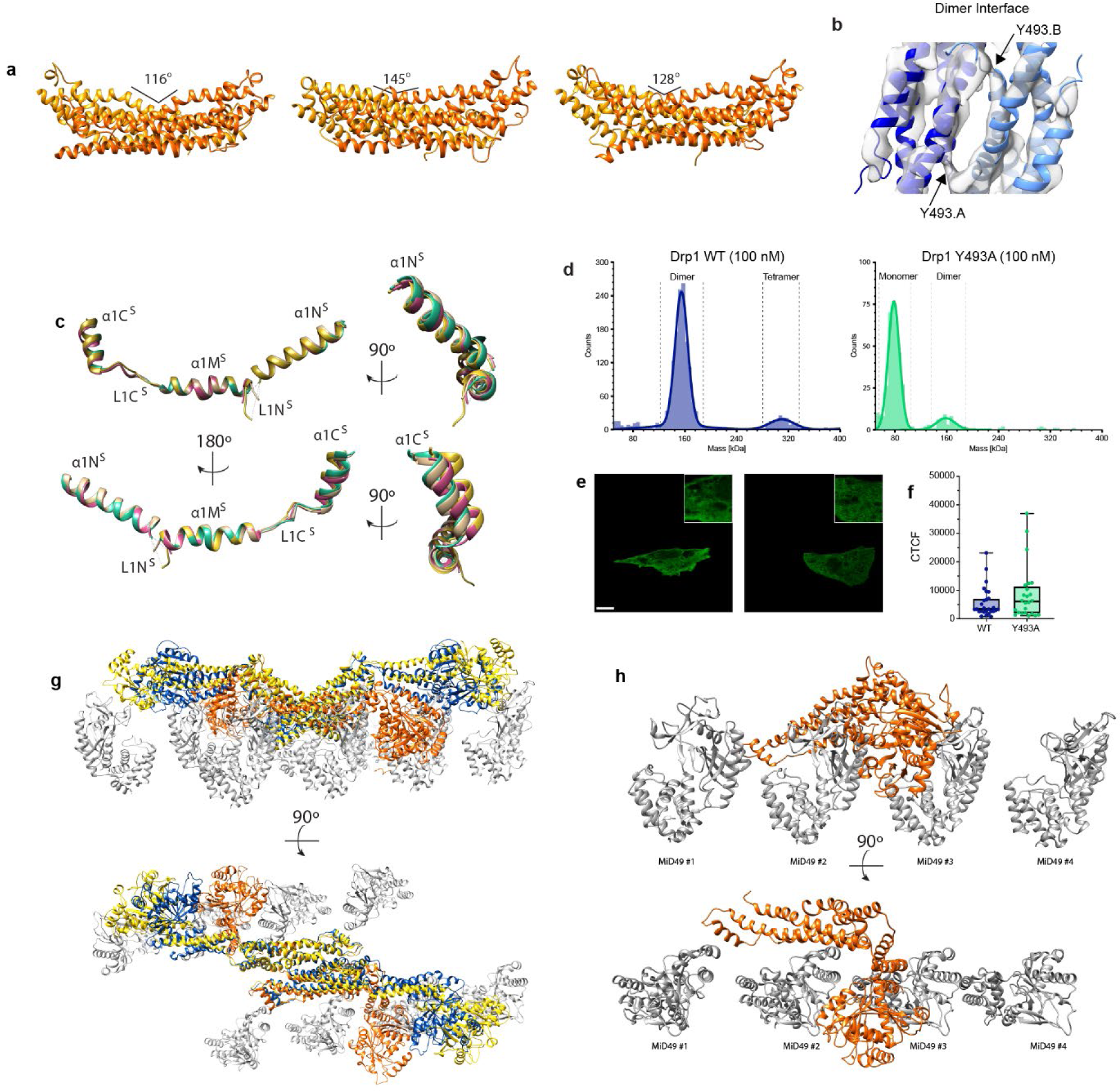
Structural Comparison of the Dimer Interface. **a**, The angle between adjacent stalks is indicated to conformations 2, 3, and 4 from single particle cryoSPARC processing and MDFF docking with AlphaFold model aligned to 4BEJ. Chain A dark orange, chain B light orange. **b**, Helices forming the dimer interface are fit within the cryo-EM density. Y493 bookends the interface. **c**, α1 helices from the crystal structure were aligned to α1Ms. (Chain A tan, chain B green, chain C pink, chain D yellow). **d**, The oligomer state of Drp1 WT and Y493A were evaluated using mass photometry at 100nM concentrations. **e**, Myc-tagged Drp1 in transfected Drp1 KO MEF cells. *Scale bar 5 µm, inset 5 µm.* **f**, Corrected Total Cell Fluorescence (CTCF) quantification for transfected cells shown in fig. 3. **g**, Alignment of Drp1 native cryo-EM structure (orange), Drp1 crystal structure (4BEJ/AlphaFold, blue), and Drp1 cryo-EM filament with MID49 and GMP-PCP (5WP9, yellow). MiD49 (gray) interfaces from 5WP9 are shown. Alignment is to reference chain A of 5WP9. **h**, Four MiD49 (gray) interfaces of 5WP9 compared to the native cryo-EM structure (orange) highlight steric clashes.

