## supplement for "Structural Basis for Regulated Assembly of the Mitochondrial Fission GTPase Drp1"

Supplement Methods

***Mass Photometry***

Mass Photometry analysis was done using a Refeyn OneMP instrument. Contrast-to-mass calibration was achieved by measuring the contrast of 4 proteins in the native marker protein standard mixture (NativeMark Unstained Protein Standard, Thermo Fisher). Four contrast values were used to generate a standard calibration curve. The experiments were performed using glass coverslips, which were thoroughly washed with Milli-Q water and isopropyl alcohol. Silicone gaskets were used for sample loading. A 1 mM protein sample was diluted to 100 nM in PBS. Movies of 6000 frames were recorded at a 100 Hz framerate using AcquireMP software and a large field-of-view acquisition setting. Data was analyzed with DiscoverMP software to produce mass values for each detected particle.

**Supplementary Table 1. Cryo-EM data collection, refinement, and validation statistics**

| | Drp1 Dimer (EMDB-40967) (PDB 8T1H) |
|---|---|
| **Data collection and processing** | |
| Magnification | 130,000X |
| Voltage (kV) | 300 |
| Electron exposure (e–/Å$^2$) | 47.76 |
| Defocus range (μm) | -0.8 to -2.0 |
| Pixel size (Å) | 1.07 |
| Symmetry imposed | C1 |
| Initial particle images (no.) | 883,690 |
| Final particle images (no.) | 71,611 |
| Map resolution (Å) | 5.97 |
| FSC threshold | 0.143 |
| Map resolution range (Å) | 13.15-5.46 |
| **Refinement** | |
| Initial model used (PDB code) | 4BEJ, AlphaFold |
| Model composition | |
| Non-hydrogen atoms | 9410 |
| Protein residues | 1190 |
| Ligands | 0 |
| R.m.s. deviations | |
| Bond lengths (Å) | 0.001 |
| Bond angles (°) | 0.358 |
| Validation | |
| MolProbity score | 1.32 |
| Clashscore | 4.56 |
| Poor rotamers (%) | |
| Ramachandran plot | |
| Favored (%) | 97.55 |
| Allowed (%) | 2.45 |
| Disallowed (%) | 0.00 |

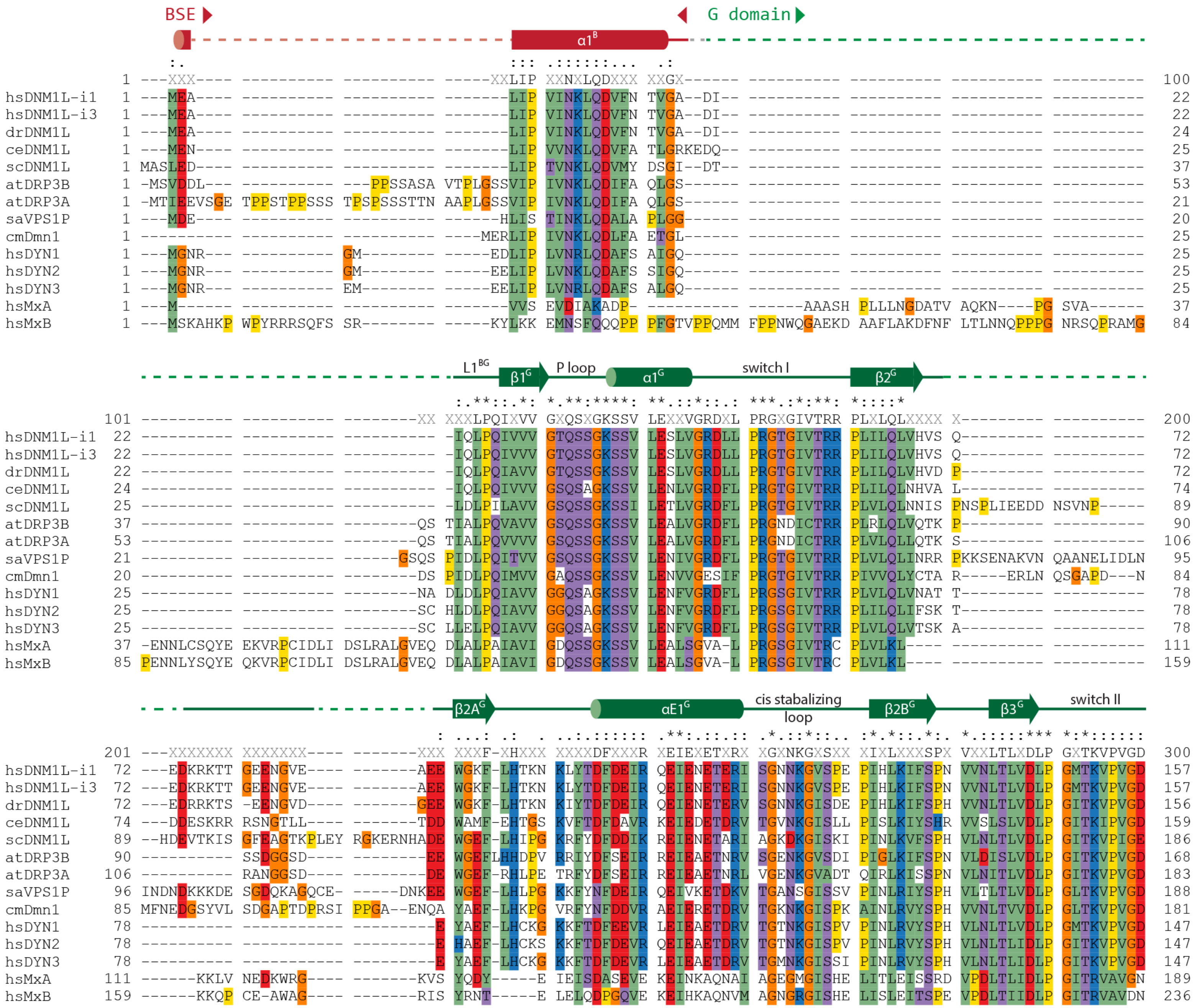
BSE
G domain
α1B
1 ---XXX---- ---------- ---------- -----XXLIP XXNXLQDXXX XXGX------ ---------- ---------- ---------- ---------- 100
hsDNM1L-i1 1 ---MEA---- ---------- ---------- -------LIP VINKLQDVFN TVGA--DI-- ---------- ---------- ---------- ---------- 22
hsDNM1L-i3 1 ---MEA---- ---------- ---------- -------LIP VINKLQDVFN TVGA--DI-- ---------- ---------- ---------- ---------- 22
drDNM1L 1 ---MEA---- ---------- ---------- -------LIP VINKLQDVFN TVGA--DI-- ---------- ---------- ---------- ---------- 24
ceDNM1L 1 ---MEN---- ---------- ---------- -------LIP VVNKLQDVFA TLGRKEDQ-- ---------- ---------- ---------- ---------- 25
scDNM1L 1 MASLED---- ---------- ---------- -------LIP TVNKLQDVMY DSGI--DT-- ---------- ---------- ---------- ---------- 37
atDRP3B 1 -MSVDDL--- ---------- ---PPSSASA VTPLGSSVIP IVNKLQDIFA QLGS------ ---------- ---------- ---------- ---------- 53
atDRP3A 1 -MTIEEVSGE TPPSTPPSSS TPSPSSSTTN AAPLGSSVIP IVNKLQDIFA QLGS------ ---------- ---------- ---------- ---------- 21
saVPS1P 1 ---MDE---- ---------- ---------- ------HLIS TINKLQDALA PLGG------ ---------- ---------- ---------- ---------- 20
cmDmn1 1 ---------- ---------- ---------- ----MERLIP IVNKLQDLFA ETGL------ ---------- ---------- ---------- ---------- 25
hsDYN1 1 ---MGNR--- ---------- GM-------- -----EDLIP LVNRLQDAFS AIGQ------ ---------- ---------- ---------- ---------- 25
hsDYN2 1 ---MGNR--- ---------- GM-------- -----EELIP LVNKLQDAFS SIGQ------ ---------- ---------- ---------- ---------- 25
hsDYN3 1 ---MGNR--- ---------- EM-------- -----EELIP LVNRLQDAFS ALGQ------ ---------- ---------- ---------- ---------- 25
hsMxA 1 ---M------ ---------- ---------- -------VVS EVDIAKADP- ---------- ------AAASH PLLLNGDATV AQKN----PG SVA------- 37
hsMxB 1 ---MSKAHKP WPYRRRSQFS SR-------- -----KYLKK EMNSFQQQPP PFGTVPPQMM FPPNWQGAEKD AAFLAKDFNF LTLNNQPPPG NRSQPRAMG 84
L1BG
β1G
P loop
α1G
switch I
β2G
101 ---------- ---------- --------XX XXXLPQIXVV GXQSXGKSSV LEXXVGRDXL PRGXGIVTRR PLXLQLXXXX X--------- ---------- 200
hsDNM1L-i1 22 ---------- ---------- ---------- -IQLPQIVVV GTQSSGKSSV LESLVGRDLL PRGTGIVTRR PLILQLVHVS Q--------- ---------- 72
hsDNM1L-i3 22 ---------- ---------- ---------- -IQLPQIVVV GTQSSGKSSV LESLVGRDLL PRGTGIVTRR PLILQLVHVS Q--------- ---------- 72
drDNM1L 22 ---------- ---------- ---------- -IQLPQIAVV GTQSSGKSSV LESLVGRDLL PRGTGIVTRR PLILQLVHVD P--------- ---------- 72
ceDNM1L 24 ---------- ---------- ---------- -IQLPQIVVV GSQSAGKSSV LENLVGRDFL PRGTGIVTRR PLILQLNHVA L--------- ---------- 74
scDNM1L 25 ---------- ---------- ---------- -LDLPILAVV GSQSSGKSSI LETLVGRDFL PRGTGIVTRR PLVLQLNNIS PNSPLIEEDD NSVNP----- 89
atDRP3B 37 ---------- ---------- --------QS TIALPQVAVV GSQSSGKSSV LEALVGRDFL PRGNDICTRR PLRLQLVQTK P--------- ---------- 90
atDRP3A 53 ---------- ---------- --------QS TIALPQVVVV GSQSSGKSSV LEALVGRDFL PRGNDICTRR PLVLQLLQTK S--------- ---------- 106
saVPS1P 21 ---------- ---------- ------GSQS PIDLPQITVV GSQSSGKSSV LENIVGRDFL PRGTGIVTRR PLVLQLINRR PKKSENAKVN QAANELIDLN 95
cmDmn1 20 ---------- ---------- --------DS PIDLPQIMVV GAQSSGKSSV LENVVGESIF PRGTGIVTRR PIVVQLYCTA R-----ERLN QSGAPD---N 84
hsDYN1 25 ---------- ---------- --------NA DLDLPQIAVV GGQSAGKSSV LENFVGRDFL PRGSGIVTRR PLVLQLVNAT T--------- ---------- 78
hsDYN2 25 ---------- ---------- --------SC HLDLPQIAVV GGQSAGKSSV LENFVGRDFL PRGSGIVTRR PLILQLIFSK T--------- ---------- 78
hsDYN3 25 ---------- ---------- --------SC LLELPQIAVV GGQSAGKSSV LENFVGRDFL PRGSGIVTRR PLVLQLVTSK A--------- ---------- 78
hsMxA 37 -ENNLCSQYE EKVRPCIDLI DSLRALGVEQ DLALPAIAVI GDQSSGKSSV LEALSGVA-L PRGSGIVTRC PLVLKL---- ---------- ---------- 111
hsMxB 85 PENNLYSQYE QKVRPCIDLI DSLRALGVEQ DLALPAIAVI GDQSSGKSSV LEALSGVA-L PRGSGIVTRC PLVLKL---- ---------- ---------- 159
β2AG
αE1G
cis stabalizing loop
β2BG
β3G
switch II
201 ---XXXXXXX XXXXXXX---- -------XXX XXXF-XHXXX XXXXDFXXXR XEIEXETXRX XGXNKGXSXX XIXLXXXSPX VXXLTLXDLP GXTKVPVGD 300
hsDNM1L-i1 72 ---EDKRKTT GEENGVE---- -------AEE WGKF-LHTKN KLYTDFDEIR QEIENETERI SGNNKGVSPE PIHLKIFSPN VVNLTLVDLP GMTKVPVGD 157
hsDNM1L-i3 72 ---EDKRKTT GEENGVE---- -------AEE WGKF-LHTKN KLYTDFDEIR QEIENETERI SGNNKGVSPE PIHLKIFSPN VVNLTLVDLP GMTKVPVGD 157
drDNM1L 72 ---EDRRKTS -EENGVD---- -------GEE WGKF-LHTKN KIYTDFDEIR QEIENETERV SGNNKGISDE PIHLKIFSPH VVNLTLVDLP GITKVPVGD 156
ceDNM1L 74 ---DDESKRR RSNGTLL---- -------TDD WAMF-EHTGS KVFTDFDAVR KEIEDETDRV TGVNKGISLL PISLKIYSHR VVSLSLVDLP GITKIPVGD 159
scDNM1L 89 --HDEVTKIS GFEAGTKPLEY RGKERNHADE WGEF-LHIPG KRFYDFDDIK REIENETARI AGKDKGISKI PINLKVFSPH VLNLTLVDLP GITKVPIGE 186
atDRP3B 90 ---------- SSDGGSD---- -------EE WGEFLHHDPV RRIYDFSEIR REIEAETNRV SGENKGVSDI PIGLKIFSPN VLDISLVDLP GITKVPVGD 168
atDRP3A 106 ---------- RANGGSD---- --------DE WGEF-RHLPE TRFYDFSEIR REIEAETNRL VGENKGVADT QIRLKISSPN VLNITLVDLP GITKVPVGD 183
saVPS1P 96 INDNDKKKDE SGDQKAGQCE- -----DNKEE WGEF-LHLPG KKFYNFDEIR QEIVKETDKV TGANSGISSV PINLRIYSPH VLTLTLVDLP GLTKVPVGD 188
cmDmn1 85 MFNEDGSYVL SDGAPTDPRSI PPGA--ENQA YAEF-LHKPG VRFYNFDDVR AEIERETDRV TGKNKGISPK AINLRVYSPH VVNLTVVDLP GLTKVPVGD 181
hsDYN1 78 ---------- ---------- ---------E YAEF-LHCKG KKFTDFEEVR LEIEAETDRV TGTNKGISPV PINLRVYSPH VLNLTLVDLP GMTKVPVGD 147
hsDYN2 78 ---------- ---------- ---------E HAEF-LHCKS KKFTDFDEVR QEIEAETDRV TGTNKGISPV PINLRVYSPH VLNLTLIDLP GITKVPVGD 147
hsDYN3 78 ---------- ---------- ---------E YAEF-LHCKG KKFTDFDEVR LEIEAETDRV TGMNKGISSI PINLRVYSPH VLNLTLIDLP GITKVPVGD 147
hsMxA 111 ------KKLV NEDKWRG---- -------KVS YQDY-----E IEISDASEVE KEINKAQNAI AGEGMGISHE LITLEISSRD VPDLTLIDLP GITRVAVGN 189
hsMxB 159 ------KKQP CE-AWAG---- -------RIS YRNT-----E LELQDPGQVE KEIHKAQNVM AGNGRGISHE LISLEITSPE VPDLTIIDLP GITRVAVDN 236

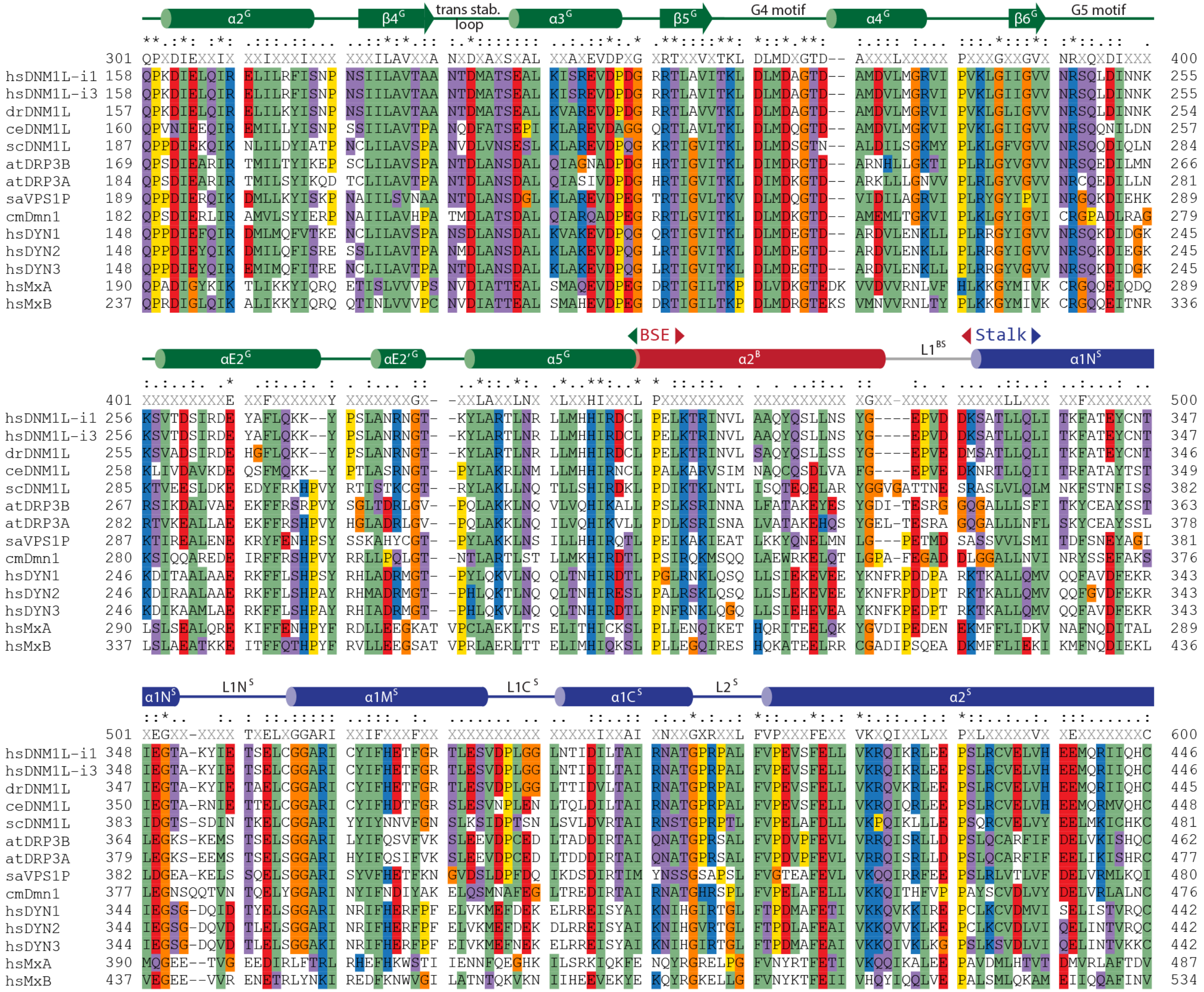
α2G
β4G
trans stab. loop
α3G
β5G
G4 motif
α4G
β6G
G5 motif
301 QPXDIEXXIX XXIXXXIXXX XXXILAVXXA NXDXAXSXAL XXAXEVDPXG XRTXXVXTKL DLMDXGTD-- AXXXLXXXXX PXXXGXXGVX NRXQXDIXXX 400
hsDNM1L-i1 158 QPKDIELQIR ELILRFISNP NSIILAVTAA NTDMATSEAL KISREVDPDG RRTLAVITKL DLMDAGTD-- AMDVLMGRVI PVKLGIIGVV NRSQLDINNK 255
hsDNM1L-i3 158 QPKDIELQIR ELILRFISNP NSIILAVTAA NTDMATSEAL KISREVDPDG RRTLAVITKL DLMDAGTD-- AMDVLMGRVI PVKLGIIGVV NRSQLDINNK 255
drDNM1L 157 QPKDIELQIR ELILKYISNP NSIILAVTAA NTDMATSEAL KVAREVDPDG RRTLAVVTKL DLMDAGTD-- AMDVLMGRVI PVKLGIIGVV NRSQLDINNK 254
ceDNM1L 160 QPVNIEEQIR EMILLYISNP SSIILAVTPA NQDFATSEPI KLAREVDAGG QRTLAVLTKL DLMDQGTD-- AMDVLMGKVI PVKLGIIGVV NRSQQNILDN 257
scDNM1L 187 QPPDIEKQIK NLILDYIATP NCLILAVSPA NVDLVNSESL KLAREVDPQG KRTIGVITKL DLMDSGTN-- ALDILSGKMY PLKLGFVGVV NRSQQDIQLN 284
atDRP3B 169 QPSDIEARIR TMILTYIKEP SCLILAVSPA NTDLANSDAL QIAGNADPDG HRTIGVITKL DIMDRGTD-- ARNHLLGKTI PLRLGYVGVV NRSQEDILMN 266
atDRP3A 184 QPSDIEARIR TMILSYIKQD TCLILAVTPA NTDLANSDAL QIASIVDPDG HRTIGVITKL DIMDKGTD-- ARKLLLGNVV PLRLGYVGVV NRCQEDILLN 281
saVPS1P 189 QPPDIERQIK DMLLKYISKP NAIILSVNAA NTDLANSDGL KLAREVDPEG TRTIGVLTKV DLMDQGTD-- VIDILAGRVI PLRYGYIPVI NRGQKDIEHK 289
cmDmn1 182 QPSDIERLIR AMVLSYIERP NAIILAVHPA TMDLATSDAL QIARQADPEG RRTLGVITKL DLMDKGTD-- AMEMLTGKVI PLKLGYIGVI CRGPADLRAG 279
hsDYN1 148 QPPDIEFQIR DMLMQFVTKE NCLILAVSPA NSDLANSDAL KVAKEVDPQG QRTIGVITKL DLMDEGTD-- ARDVLENKLL PLRRGYIGVV NRSQKDIDGK 245
hsDYN2 148 QPPDIEYQIK DMILQFISRE SSLILAVTPA NMDLANSDAL KLAKEVDPQG LRTIGVITKL DLMDEGTD-- ARDVLENKLL PLRRGYIGVV NRSQKDIEGK 245
hsDYN3 148 QPPDIEYQIR EMIMQFITRE NCLILAVTPA NTDLANSDAL KLAKEVDPQG LRTIGVITKL DLMDEGTD-- ARDVLENKLL PLRRGYVGVV NRSQKDIDGK 245
hsMxA 190 QPADIGYKIK TLIKKYIQRQ ETISLVVVPS NVDIATTEAL SMAQEVDPEG DRTIGILTKP DLVDKGTEDK VVDVVRNLVF HLKKGYMIVK CRGQQEIQDQ 289
hsMxB 237 QPRDIGLQIK ALIKKYIQRQ QTINLVVVPC NVDIATTEAL SMAHEVDPEG DRTIGILTKP DLMDRGTEKS VMNVVRNLTY PLKKGYMIVK CRGQQEITNR 336
BSE
Stalk
αE2G
αE2′G
α5G
α2B
L1BS
α1NS
401 XXXXXXXXXE XXFXXXXXXY XXXXXXXGX- -XXLAXXLNX XLXXHIXXXL PXXXXXXXXX XXXXXXXXXX XGXX-XXXXX XXXXXLLXXX XXFXXXXXXX 500
hsDNM1L-i1 256 KSVTDSIRDE YAFLQKK--Y PSLANRNGT- -KYLARTLNR LLMHHIRDCL PELKTRINVL AAQYQSLLNS YG----EPVD DKSATLLQLI TKFATEYCNT 347
hsDNM1L-i3 256 KSVTDSIRDE YAFLQKK--Y PSLANRNGT- -KYLARTLNR LLMHHIRDCL PELKTRINVL AAQYQSLLNS YG----EPVD DKSATLLQLI TKFATEYCNT 347
drDNM1L 255 KSVADSIRDE HGFLQKK--Y PSLANRNGT- -KYLARTLNR LLMHHIRDCL PELKTRINVL SAQYQSLLSS YG----EPVE DMSATLLQLI TKFATEYCNT 346
ceDNM1L 258 KLIVDAVKDE QSFMQKK--Y PTLASRNGT- -PYLAKRLNM LLMHHIRNCL PALKARVSIM NAQCQSDLVA FG----EPVE DKNRTLLQII TRFATAYTST 349
scDNM1L 285 KTVEESLDKE EDYFRKHPVY RTISTKCGT- -RYLAKLLNQ TLLSHIRDKL PDIKTKLNTL ISQTEQELAR YGGVGATTNE SRASLVLQLM NKFSTNFISS 382
atDRP3B 267 RSIKDALVAE EKFFRSRPVY SGLTDRLGV- -PQLAKKLNQ VLVQHIKALL PSLKSRINNA LFATAKEYES YGDI-TESRG GQGALLLSFI TKYCEAYSST 363
atDRP3A 282 RTVKEALLAE EKFFRSHPVY HGLADRLGV- -PQLAKKLNQ ILVQHIKVLL PDLKSRISNA LVATAKEHQS YGEL-TESRA GQGALLLNFL SKYCEAYSSL 378
saVPS1P 287 KTIREALENE KRYFENHPSY SSKAHYCGT- -PYLAKKLNS ILLHHIRQTL PEIKAKIEAT LKKYQNELMN LG---PETMD SASSVVLSMI TDFSNEYAGI 381
cmDmn1 280 KSIQQAREDE IRFFRSHPVY RRLLPQLGT- -NTLARTLST LLMKHIRDTL PSIRQKMSQQ LAEWRKELQT LGPA-FEGAD DLGGALLNVI NRYSSEFAKS 376
hsDYN1 246 KDITAALAAE RKFFLSHPSY RHLADRMGT- -PYLQKVLNQ QLTNHIRDTL PGLRNKLQSQ LLSIEKEVEE YKNFRPDDPA RKTKALLQMV QQFAVDFEKR 343
hsDYN2 246 KDIRAALAAE RKFFLSHPAY RHMADRMGT- -PHLQKTLNQ QLTNHIRESL PALRSKLQSQ LLSLEKEVEE YKNFRPDDPT RKTKALLQMV QQFGVDFEKR 343
hsDYN3 246 KDIKAAMLAE RKFFLSHPAY RHIADRMGT- -PHLQKVLNQ QLTNHIRDTL PNFRNKLQGQ LLSIEHEVEA YKNFKPEDPT RKTKALLQMV QQFAVDFEKR 343
hsMxA 290 LSLSEALQRE KIFFENHPYF RDLLEEGKAT VPCLAEKLTS ELITHICKSL PLLENQIKET HQRITEELQK YGVDIPEDEN EKMFFLIDKV NAFNQDITAL 289
hsMxB 337 LSLAEATKKE ITFFQTHPYF RVLLEEGSAT VPRLAERLTT ELIMHIQKSL PLLEGQIRES HQKATEELRR CGADIPSQEA DKMFFLIEKI KMFNQDIEKL 436
α1NS
L1NS
α1MS
L1CS
α1CS
L2S
α2S
501 XEGXX-XXXX TXELXGGARI XXIFXXXFXX XXXXXXXXXX XXXXXIXXAI XNXXGXRXXL FVPXXXFEXX VKXQIXXLXX PXLXXXXXVX XEXXXXXXXC 600
hsDNM1L-i1 348 IEGTA-KYIE TSELCGGARI CYIFHETFGR TLESVDPLGG LNTIDILTAI RNATGPRPAL FVPEVSFELL VKRQIKRLEE PSLRCVELVH EEMQRIIQHC 446
hsDNM1L-i3 348 IEGTA-KYIE TSELCGGARI CYIFHETFGR TLESVDPLGG LNTIDILTAI RNATGPRPAL FVPEVSFELL VKRQIKRLEE PSLRCVELVH EEMQRIIQHC 446
drDNM1L 347 IEGTA-KYIE TAELCGGARI CYIFHETFGR TLESVDPLGG LTTIDVLTAI RNATGPRPAL FVPEVSFELL VKRQVKRLEE PSLRCVELVH EEMQRIIQHC 445
ceDNM1L 350 IEGTA-RNIE TTELCGGARI CYIFHDTFGR SLESVNPLEN LTQLDILTAI RNATGPRPAL FVPEVSFELL VKRQIQRLEE PSLRCVELVH EEMQRMVQHC 448
scDNM1L 383 IDGTS-SDIN TKELCGGARI YYIYNNVFGN SLKSIDPTSN LSVLDVRTAI RNSTGPRPTL FVPELAFDLL VKPQIKLLLE PSQRCVELVY EELMKICHKC 481
atDRP3B 364 LEGKS-KEMS TSELSGGARI LYIFQSVFVK SLEEVDPCED LTADDIRTAI QNATGPRSAL FVPDVPFEVL VRRQISRLLD PSLQCARFIF DELVKISHQC 462
atDRP3A 379 LEGKS-EEMS TSELSGGARI HYIFQSIFVK SLEEVDPCED LTDDDIRTAI QNATGPRSAL FVPDVPFEVL VRRQISRLLD PSLQCARFIF EELIKISHRC 477
saVPS1P 382 LDGEA-KELS SQELSGGARI SYVFHETFKN GVDSLDPFDQ IKDSDIRTIM YNSSGSAPSL FVGTEAFEVL VKQQIRRFEE PSLRLVTLVF DELVRMLKQI 480
cmDmn1 377 LEGNSQQTVN TQELYGGARI NYIFNDIYAK ELQSMNAFEG LTREDIRTAI RNATGHRSPL FVPELAFELL VKKQITHFVP PAYSCVDLVY DELVRLALNC 476
hsDYN1 344 IEGSG-DQID TYELSGGARI NRIFHERFPF ELVKMEFDEK ELRREISYAI KNIHGIRTGL FTPDMAFETI VKKQVKKIRE PCLKCVDMVI SELISTVRQC 442
hsDYN2 344 IEGSG-DQVD TLELSGGARI NRIFHERFPF ELVKMEFDEK DLRREISYAI KNIHGVRTGL FTPDLAFEAI VKKQVVKLKE PCLKCVDLVI QELINTVRQC 442
hsDYN3 344 IEGSG-DQVD TLELSGGAKI NRIFHERFPF EIVKMEFNEK ELRREISYAI KNIHGIRTGL FTPDMAFEAI VKKQIVKLKG PSLKSVDLVI QELINTVKKC 442
hsMxA 390 MQGEE--TVG EEDIRLFTRL RHEFHKWSTI IENNFQEGHK ILSRKIQKFE NQYRGRELPG FVNYRTFETI VKQQIKALEE PAVDMLHTVT DMVRLAFTDV 487
hsMxB 437 VEGEE--VVR ENETRLYNKI REDFKNWVGI LATNTQKVKN IIHEEVEKYE KQYRGKELLG FVNYKTFEII VHQYIQQLVE PALSMLQKAM EIIQQAFINV 534

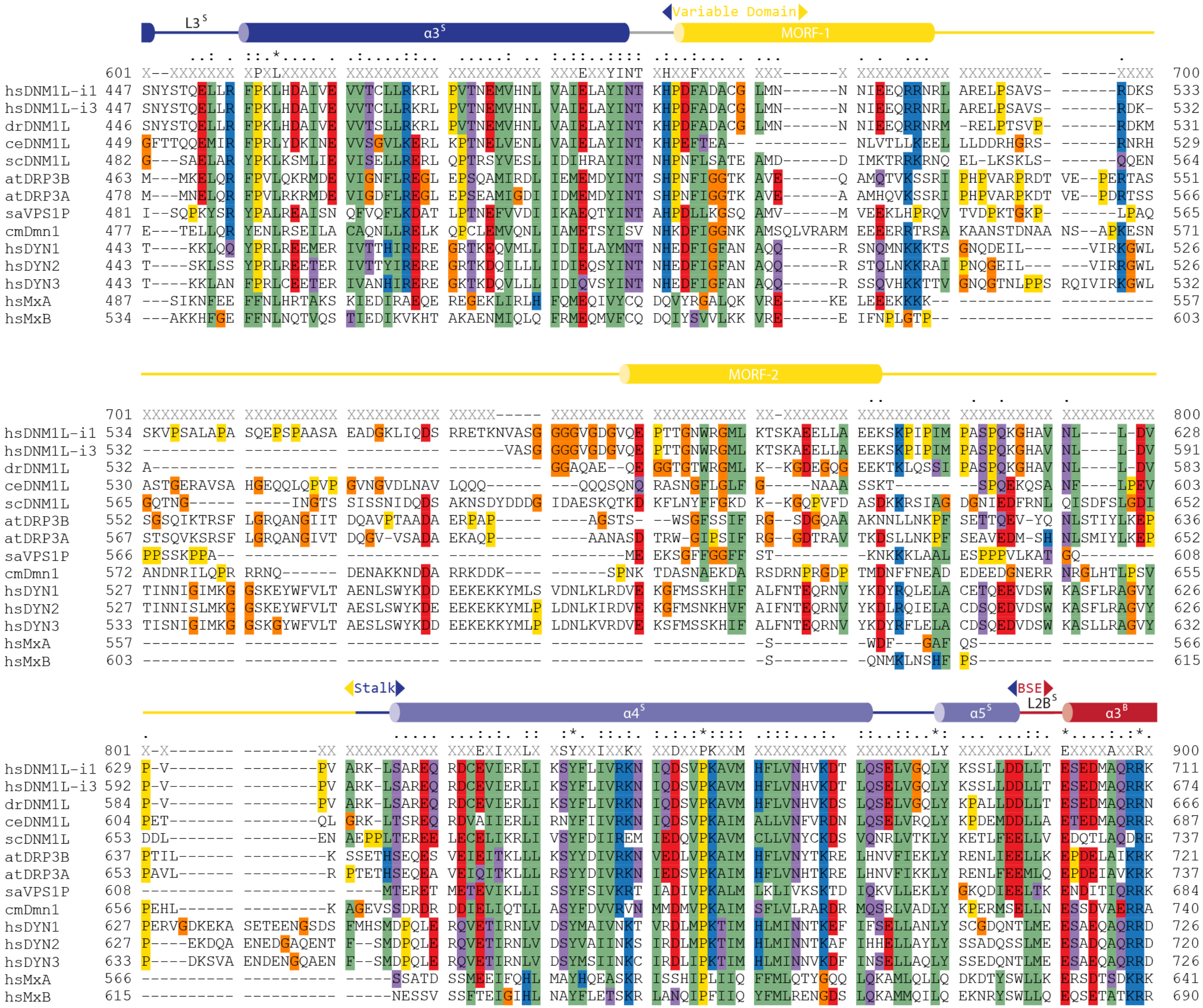
L3S
α3S
Variable Domain
MORF-1
601 X--XXXXXXX XPXLXXXXXX XXXXXXXXXX XXXXXXXXXX XXXEXXYINT XHXXFXXXXX XXX------X XXXXXXXXXX XXXXXXXXX- ------XXXX 700
hsDNM1L-i1 447 SNYSTQELLR FPKLHDAIVE VVTCLLRKRL PVTNEMVHNL VAIELAYINT KHPDFADACG LMN------N NIEEQRRNRL ARELPSAVS- ------RDKS 533
hsDNM1L-i3 447 SNYSTQELLR FPKLHDAIVE VVTCLLRKRL PVTNEMVHNL VAIELAYINT KHPDFADACG LMN------N NIEEQRRNRL ARELPSAVS- ------RDK- 532
drDNM1L 446 SNYSTQELLR FPKLHDAIVE VVTSLLRKRL PVTNEMVHNL VAIELAYINT KHPDFADACG LMN------N NIEEQRRNRM -RELPTSVP- ------RDKM 531
ceDNM1L 449 GFTTQQEMIR FPRLYDKINE VVSGVLKERL KPTNELVENL VAIELAYINT KHPEFTEA-- ---------- NLVTLLKEEL LLDDRHGRS- ------RNRH 529
scDNM1L 482 G---SAELAR YPKLKSMLIE VISELLRERL QPTRSYVESL IDIHRAYINT NHPNFLSATE AMD------D IMKTRRKRNQ EL-LKSKLS- ------QQEN 564
atDRP3B 463 M---MKELQR FPVLQKRMDE VIGNFLREGL EPSQAMIRDL IEMEMDYINT SHPNFIGGTK AVE------Q AMQTVKSSRI PHPVARPRDT VE--PERTAS 551
atDRP3A 478 M---MNELQR FPVLRKRMDE VIGDFLREGL EPSEAMIGDI IDMEMDYINT SHPNFIGGTK AVE------A AMHQVKSSRI PHPVARPKDT VE--PDRTSS 566
saVPS1P 481 I--SQPKYSR YPALREAISN QFVQFLKDAT LPTNEFVVDI IKAEQTYINT AHPDLLKGSQ AMV------M VEEKLHPRQV TVDPKTGKP- ------LPAQ 565
cmDmn1 477 E---TELLQR YENLRSEILA CAQNLLRELK QPCLEMVQNL IAMETSYISV NHKDFIGGNK AMSQLVRARM EEEERRTRSA KAANSTDNAA NS--APKESN 571
hsDYN1 443 T----KKLQQ YPRLREEMER IVTTHIRERE GRTKEQVMLL IDIELAYMNT NHEDFIGFAN AQQ------R SNQMNKKKTS GNQDEIL--- ---VIRKGWL 526
hsDYN2 443 T----SKLSS YPRLREETER IVTTYIRERE GRTKDQILLL IDIEQSYINT NHEDFIGFAN AQQ------R STQLNKKRAI PNQGEIL--- ---VIRRGWL 526
hsDYN3 443 T----KKLAN FPRLCEETER IVANHIRERE GKTKDQVLLL IDIQVSYINT NHEDFIGFAN AQQ------R SSQVHKKTTV GNQGTNLPPS RQIVIRKGWL 532
hsMxA 487 ---SIKNFEE FFNLHRTAKS KIEDIRAEQE REGEKLIRLH FQMEQIVYCQ DQVYRGALQK VRE-----KE LEEEKKKK-- ---------- ---------- 557
hsMxB 534 ---AKKHFGE FFNLNQTVQS TIEDIKVKHT AKAENMIQLQ FRMEQMVFCQ DQIYSVVLKK VRE------E IFNPLGTP-- ---------- ---------- 603
MORF-2
701 XXXXXXXXXX XXXXXXXXXX XXXXXXXXXX XXXXX----- XXXXXXXXXX XXXXXXXXXX XX-XXXXXXX XXXXXXXXXX XXXXXXXXXX XXXXXXXXXX 800
hsDNM1L-i1 534 SKVPSALAPA SQEPSPAASA EADGKLIQDS RRETKNVASG GGGVGDGVQE PTTGNWRGML KTSKAEELLA EEKSKPIPIM PASPQKGHAV NL----L-DV 628
hsDNM1L-i3 532 ---------- ---------- ---------- ------VASG GGGVGDGVQE PTTGNWRGML KTSKAEELLA EEKSKPIPIM PASPQKGHAV NL----L-DV 591
drDNM1L 532 A--------- ---------- ---------- ---------- GGAQAE--QE GGTGTWRGML K--KGDEGQG EEKTKLQSSI PASPQKGHAV NL----L-DV 583
ceDNM1L 530 ASTGERAVSA HGEQQLQPVP GVNGVDLNAV LQQQ------ ---QQQSQNQ RASNGFLGLF G-----NAAA SSKT------ --SPQEKQSA NF----LPEV 603
scDNM1L 565 GQTNG----- -----INGTS SISSNIDQDS AKNSDYDDDG IDAESKQTKD KFLNYFFGKD K--KGQPVFD ASDKKRSIAG DGNIEDFRNL QISDFSLGDI 652
atDRP3B 552 SGSQIKTRSF LGRQANGIIT DQAVPTAADA ERPAP----- ----AGSTS- --WSGFSSIF RG--SDGQAA AKNNLLNKPF SETTQEV-YQ NLSTIYLKEP 636
atDRP3A 567 STSQVKSRSF LGRQANGIVT DQGV-VSADA EKAQP----- ----AANASD TRW-GIPSIF RG--GDTRAV TKDSLLNKPF SEAVEDM-SH NLSMIYLKEP 652
saVPS1P 566 PPSSKPPA-- ---------- ---------- ---------- --------ME EKSGFFGGFF ST-------- -KNKKKLAAL ESPPPVLKAT GQ-------- 608
cmDmn1 572 ANDNRILQPR RRNQ------ DENAKKNDDA RRKDDK---- ------SPNK TDASNAEKDA RSDRNPRGDP TMDNFFNEAD EDEEDGNERR NRGLHTLPSV 655
hsDYN1 527 TINNIGIMKG GSKEYWFVLT AENLSWYKDD EEKEKKYMLS VDNLKLRDVE KGFMSSKHIF ALFNTEQRNV YKDYRQLELA CETQEEVDSW KASFLRAGVY 626
hsDYN2 527 TINNISLMKG GSKEYWFVLT AESLSWYKDE EEKEKKYMLP LDNLKIRDVE KGFMSNKHVF AIFNTEQRNV YKDLRQIELA CDSQEDVDSW KASFLRAGVY 626
hsDYN3 533 TISNIGIMKG GSKGYWFVLT AESLSWYKDD EEKEKKYMLP LDNLKVRDVE KSFMSSKHIF ALFNTEQRNV YKDYRFLELA CDSQEDVDSW KASLLRAGVY 632
hsMxA 557 ---------- ---------- ---------- ---------- ---------- ---------- -S-------- -WDF---GAF QS-------- ---------- 566
hsMxB 603 ---------- ---------- ---------- ---------- ---------- ---------- -S-------- -QNMKLNSHF PS-------- ---------- 615
Stalk
α4S
BSE
α5S
L2BS
α3B
801 X-X------- --------XX XXXXXXXXXX XXXEXIXXLX XSYXXIXXKX XXDXXPKXXM XXXXXXXXXX XXXXXXXXLY XXXXXXXLXX EXXXXAXXRX 900
hsDNM1L-i1 629 P-V------- --------PV ARK-LSAREQ RDCEVIERLI KSYFLIVRKN IQDSVPKAVM HFLVNHVKDT LQSELVGQLY KSSLLDDLLT ESEDMAQRRK 711
hsDNM1L-i3 592 P-V------- --------PV ARK-LSAREQ RDCEVIERLI KSYFLIVRKN IQDSVPKAVM HFLVNHVKDT LQSELVGQLY KSSLLDDLLT ESEDMAQRRK 674
drDNM1L 584 P-V------- --------PV ARK-LSAREQ RDCEVIERLI KSYFLIVRKN IQDSVPKAVM HFLVNHVKDS LQSELVGQLY KPALLDDLLT ESEDMAQRRN 666
ceDNM1L 604 PET------- --------QL GRK-LTSREQ RDVAIIERLI RNYFIIVRKN IQDSVPKAIM ALLVNFVRDN LQSELVRQLY KPDEMDDLLA ETEDMAQRRR 687
scDNM1L 653 DDL------- --------EN AEPPLTEREE LECELIKRLI VSYFDIIREM IEDQVPKAVM CLLVNYCKDS VQNRLVTKLY KETLFEELLV EDQTLAQDRE 737
atDRP3B 637 PTIL------ ---------K SSETHSEQES VEIEITKLLL KSYYDIVRKN VEDLVPKAIM HFLVNYTKRE LHNVFIEKLY RENLIEELLK EPDELAIKRK 721
atDRP3A 653 PAVL------ ---------R PTETHSEQEA VEIQITKLLL RSYYDIVRKN IEDSVPKAIM HFLVNHTKRE LHNVFIKKLY RENLFEEMLQ EPDEIAVKRK 737
saVPS1P 608 ---------- ---------- ----MTERET METEVIKLLI SSYFSIVKRT IADIVPKALM LKLIVKSKTD IQKVLLEKLY GKQDIEELTK ENDITIQRRK 684
cmDmn1 656 PEHL------ ---------K AGEVSSDRDR DDIELIQTLL ASYFDVVRVN MMDMVPKAIM SFLVLRARDR MQSRLVADLY KPERMSELLN ESSDVAERRA 740
hsDYN1 627 PERVGDKEKA SETEENGSDS FMHSMDPQLE RQVETIRNLV DSYMAIVNKT VRDLMPKTIM HLMINNTKEF IFSELLANLY SCGDQNTLME ESAEQAQRRD 726
hsDYN2 627 P----EKDQA ENEDGAQENT F--SMDPQLE RQVETIRNLV DSYVAIINKS IRDLMPKTIM HLMINNTKAF IHHELLAYLY SSADQSSLME ESADQAQRRD 720
hsDYN3 633 P----DKSVA ENDENGQAEN F--SMDPQLE RQVETIRNLV DSYMSIINKC IRDLIPKTIM HLMINNVKDF INSELLAQLY SSEDQNTLME ESAEQAQRRD 726
hsMxA 566 ---------- ---------- -----SSATD SSMEEIFQHL MAYHQEASKR ISSHIPLIIQ FFMLQTYGQQ LQKAMLQLLQ DKDTYSWLLK ERSDTSDKRK 641
hsMxB 615 ---------- ---------- -----NESSV SSFTEIGIHL NAYFLETSKR LANQIPFIIQ YFMLRENGDS LQKAMMQILQ EKNRYSWLLQ EQSETATKRR 690

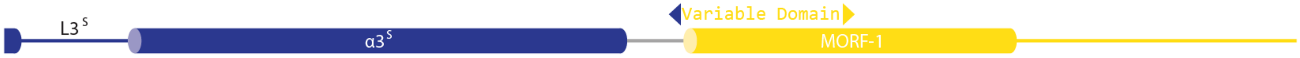

```
                   .......:.  .:  ..:...
             901 XXXXXXXXLX XAXXXXXXXX XXXXX----- ---------- ---------- ---------- ---------- ---------- ---------- ---------- 1000
hsDNM1L-i1   712 EAADMLKALQ GASQIIAEIR ETHLW----- ---------- ---------- ---------- ---------- ---------- ---------- ----------  736
hsDNM1L-i3   675 EAADMLKALQ GASQIIAEIR ETHLW----- ---------- ---------- ---------- ---------- ---------- ---------- ----------  699
drDNM1L      667 EAADMLKALQ KASQVIAEIR ETHLW----- ---------- ---------- ---------- ---------- ---------- ---------- ----------  691
ceDNM1L      688 DTLETMKALQ QASVIISEVR ETQVW----- ---------- ---------- ---------- ---------- ---------- ---------- ----------  712
scDNM1L      738 LCVKSLGVYK KAATLISNIL ---------- ---------- ---------- ---------- ---------- ---------- ---------- ----------  757
atDRP3B      722 RTQETLRILQ QANRTLDELP LEAESVERGY ---------- KI-GSEAKHE ELPGT-RRSR T----ETNGN GRLHM----- ---------- ----------  780
atDRP3A      738 RTQETLHVLQ QAYRTLDELP LEADSVSAGM SKHQELLTSS KY-STSSSYS ASPSTTRRSR RAGDQHQNGY GF-------- ---------- ----------  808
saVPS1P      685 ECKKMVEILR NASQIVSTV- ---------- ---------- ---------- ---------- ---------- ---------- ---------- ----------  703
cmDmn1       741 TAKRMVDLLQ RGMAVINEVR DVHVKEEI-- ---------- ---------- --------------------- ---------- ---------- ---------  768
hsDYN1       727 EMLRMYHALK EALSIIGDIN TTTVSTPMPP PVDDSWLQVQ SVPAGRRSPT SSPTPQRRAP -AV-PPARPG S-RGPAPGPP ----PAGSAL G-GAPPVPSR  818
hsDYN2       721 DMLRMYHALK EALNIIGDIS TSTVSTPVPP PVDDTWLQSA S--------S HSPTPQRRPV SSIHPPGRPP AVRGPTPGPP LIPVPVGAAA SFSAPPIPSR  812
hsDYN3       727 EMLRMYQALK EALGIIGDIS TATVSTPAPP PVDDSWIQ-- ---HSRRSPP PSPTTQRRPT LSA-PLARPT SGRGPAPAIP ----SPGPH- S-GAPPVPFR  814
hsMxA        642 FLKERLARLT QARRRLAQFP G--------- ---------- ---------- ---------- ---------- ---------- ---------- ----------  662
hsMxB        691 ILKERIYRLT QARHALCQFS SKEIH----- ---------- ---------- ---------- ---------- ---------- ---------- ----------  715

            1001 ---------- ---------- ---------- ---------- ---------- ---------  1100
hsDNM1L-i1   736 ---------- ---------- ---------- ---------- ---------- ---------   736
hsDNM1L-i3   699 ---------- ---------- ---------- ---------- ---------- ---------   699
drDNM1L      691 ---------- ---------- ---------- ---------- ---------- ---------   691
ceDNM1L      712 ---------- ---------- ---------- ---------- ---------- ---------   712
scDNM1L      757 ---------- ---------- ---------- ---------- ---------- ---------   757
atDRP3B      780 ---------- ---------- ---------- ---------- ---------- ---------   780
atDRP3A      808 ---------- ---------- ---------- ---------- ---------- ---------   808
saVPS1P      703 ---------- ---------- ---------- ---------- ---------- ---------   703
cmDmn1       768 ---------- ---------- ---------- ---------- ---------- ---------   768
hsDYN1       819 PGASP----- -DPFGPPPQV PSRPNRAPPG ----VPSRSG QASPSRPESP RPPFDL---   864
hsDYN2       813 PGPQSVF-AN SDLFPAPPQI PSRPVRIPPG IPPGVPSRRP PAAPSRPTII RPAEPSLLD   870
hsDYN3       815 PGPLPPFPSS SDSFGAPPQV PSRPTRAPPS ----VPSRRP PPSPTRPTII RPLESSLLD   869
hsMxA        662 ---------- ---------- ---------- ---------- ---------- ---------   662
hsMxB        715 ---------- ---------- ---------- ---------- ---------- ---------   715
```

**Supplementary Data 1. | Full Sequence Alignment of Selected DSPs.** Amino acid sequences of the two splice variants of *Homo sapiens* (hs) DNM1L isoforms (Swiss-Prot accession O00429-1,3), *Danio rerio* (dr) DNM1L (Q7SXN5), *Caenorhabditis elegans* (ce) DNM1L (Q8WQC9), *Saccharomyces cerevisiae* (sc) DNM1L (P54861), *Arabidopsis thaliana* (at) DRP3B (Q8LFT2), *Arabidopsis thaliana* (at) DRP3A (Q8S944), *Saccharomyces arboricola* (sa) VPS1P (J8LLG7), *Cyanidioschyzon merolae* (cm) DMN1 (Q84Y91), *Homo sapiens* DYN1 (Q05193), *Homo sapiens* DYN2 (P50570), *Homo sapiens* DYN3 (Q9UQ16), *Homo sapiens* MxA (P20591) and *Homo sapiens* (P20592) were aligned using Clustal X in SnapGene and color classifications were manually adjusted. Positively charged residues are blue (D,E), negatively charged residues are red (R, H, K), polar residues are purple (S, T, N, Q), hydrophobic residues are green (A, V, I, L, M, F, Y, W), prolines are yellow, and glycines are orange. A "-" signifies no sequence or >15% consensus, a "X" signifies any amino acid is present at least 15% frequency but with no consensus, a "." Signified >35% and <75% consensus, a ":" signifies a >75% and <100% consensus, and a "*" signifies a 100% consensus.
